# Leaf bacterial community structure and variation in wild ruderal plants are shaped by the interaction of host species and defense chemistry with environment

**DOI:** 10.1101/2022.03.16.484556

**Authors:** Teresa Mayer, Michael Reichelt, Jonathan Gershenzon, Matthew Agler

## Abstract

Variable phenotypes help plants ensure fitness and survival in the face of unpredictable environmental stresses. Leaf bacteria (bacteriomes) can extend plant phenotypes and are well-known to vary from one plant to the next, but little is known about controls on this variation. Here, we find in 9 populations of *Arabidopsis thaliana* that core leaf bacteriomes are largely, but not completely, shared with other ground-dwelling ruderal plant species. Strength of differentiation between plant species and between *A. thaliana* populations shifts from year to year, becoming stronger when plants within populations have more similar leaf bacteriomes (less plant-to-plant variation or stochasticity). Interestingly, across different populations, plants with shared leaf aliphatic glucosinolate chemotypes exhibited similar strong year-to-year stochasticity shifts. Therefore, stochasticity of leaf bacteriomes in plant populations changes in specific ways and might be controlled by plant traits, with important implications for how plants adapt to complex and shifting environments.

## Introduction

Plants are colonized by diverse organisms including bacteria, fungi and protists. These microorganisms not only use plants as a primary resource but can provide important functions in return. Besides well-known beneficial fungal-plant relationships, such as arbuscular mycorrhiza, other microorganisms like bacteria can play critical roles in plant survival. For example, bacteria are required for plants to survive detrimental effects of normal soil fungi and oomycetes (Duran et al., 2018). On leaves, bacteria can function for example as barriers to the constant threat posed by pathogen invasion (Berg and Koskella, 2018; Ritpitakphong et al., 2016). Thus, much effort is put into understanding how leaf bacterial communities (bacteriomes) are shaped to help develop new, sustainable approaches to agriculture and biodiversity management.

Compared to other organs, leaf bacteriomes are still relatively poorly understood. Different plant species and plant genotypes tend to have overlapping leaf microbiota on the phylum level with differences at finer levels (Knief et al., 2010; Massoni et al., 2020). Similar to rhizosphere colonizers, the primary source of the leaf microbiota is thought to be soil (Massoni et al., 2021; Tkacz et al., 2020), but vertical colonization and deposition from air could also play important roles (García-Suárez et al., 2017; Maignien et al., 2014). Selection of bacterial plant colonizers from complex inocula is influenced to some extent by factors like host genetics and host age (Bodenhausen et al., 2014; Wagner et al., 2016) and much of this selection may happen early in the plant life cycle since early colonizers seem to be very robust to later invasion (Carlstrom et al., 2019; Leopold and Busby, 2020). In nature, however, leaf microbiomes are not necessarily completely stable and shifts across the life cycle of plants are well-documented (Williams et al., 2013). Indeed, leaves are highly dynamic, exposed environments and both abiotic factors like temperature, humidity and UV (Aydogan et al., 2020) and biological factors like pathogens can strongly shape communities (Agler et al., 2016). Hosts can also strongly interact with these factors, resulting in strong interaction effects between plant host factors and environments on leaf microbiome structures (Wagner et al., 2016).

Some threats to plants that can be mitigated with help from the microbiome are likely to be endemic, specific to the locations and soils where the plants grow (Agler et al., 2016). On the other hand, many selective pressures on a given plant population are not necessarily constant and in nature can vary in magnitude and even direction, remaining unpredictable from one year to the next (Postma and Ågren, 2018). Leaf bacteria can help plants to adapt and thrive in complex environments not just by improving certain plant phenotypes but also by making phenotypes more plastic and variable (Hawkes et al., 2021; Petipas et al., 2021). Therefore, plants should be expected not only to structure leaf bacteriomes in specific ways to address certain threats, but also to manage community variation to help ensure survival in the face of unpredictability. Consistent with this, we observed that *A. thaliana* leaf bacteriomes planted in three years in a common garden consistently converged toward a specific community structure, but with significant plant-to-plant variation (Almario et al., 2021). An important but largely unaddressed question, then, is not just how leaf bacteriomes are shaped into specific structures, but also how *variation* arises and what factors might control it.

Some of these questions about how microorganisms and plants collaborate to adaptat to complex, changing natural environments can be addressed by studying wild plant populations. Persistent and numerous wild populations of *A. thaliana* can be found throughout Europe (Thiergart et al., 2020). In much of Europe, *A. thaliana* populations grow together with other diverse ruderal species and *A. thaliana* itself can have a high trait diversity even on small spatial scales (Katz et al., 2021); this diversity is so far underutilized in microbiome research but can be leveraged to understand how host factors shape colonization in nature. In this study, we begin to address some of these questions by looking into 13 distinct *A. thaliana* populations in Jena, Germany, all within a 4km radius. By sampling whole leaf bacteriomes of both *A. thaliana* and other sympatric plants across multiple years, we asked how they differ between locations and between host species and whether these effects are consistent across time. We identify stochasticity of leaf bacteriomes as an important aspect defining whether leaf bacteriomes are distinguishable between host species or between the populations. Interestingly, this stochasticity changes between populations and over years and is potentially linked to *A. thaliana* leaf chemotypes.

## Material & Methods

### Isolation and characterization of wild *A. thaliana* population lines

#### Overview

In spring 2018, we identified 13 wild *A. thaliana* populations in Jena, Germany (**Figure S1A**). Plants in the populations generally occupy similar niches – growth in sandy soil around or between cobblestones or sidewalks. We collected leaf samples of nine plants growing naturally in each local population to extract DNA to infer genotypic heterogeneity using microsatellite length polymorphisms. Additionally, around five plants were collected from each population, re-potted and grown in the lab to collect seeds. For isolated lines of seven of the populations, we characterized leaf glucosinolate (GL) profiles and later confirmed them in a second batch after plants had been selfed to generate mostly homozygous lines. Additionally, we selected five populations (see above) with completely uniform microsatellite profiles (NG2 - Neugasse2, PB - Paradiesbahnhof, SW1 - Sandweg1, JT1 - Johannistor1, Woe - Wöllnitzer) and characterized plant development times in the selfed lines. Seeds of four of the populations were donated to the NASC seed database and given new names; seeds of the population PB will donated as soon as possible. An overview of the data available and NASC accession numbers for each population is provided in **Table S2**.

#### Plant DNA extraction of leaf samples

We rapidly extracted DNA of the nine plants from each local plant population as well as the control lines Col-0 and Ws-0 (Edwards et al., 1991). We crushed one leaf with a metal pestle, added 400 μL extraction buffer (200 mM Tris HCl pH 8.0, 250 mM NaCl, 25 mM EDTA, 0.5% SDS) and vortexed for 5 s. We centrifuged the samples at 19,000 x g for 1 min, transferred the supernatant to a fresh tube and added 300 μL of isopropanol. Samples were incubated at RT for 2 min. We centrifuged the samples again and removed the supernatant. After the DNA was air-dried, we dissolved the pellets in 100 μL TE-Buffer (10 mM Tris-HCl pH 8.0, 1 mM EDTA pH 8.0).

#### Microsatellite diversity

We assessed the intra- and interspecies diversity of the plant population by the diversity of three microsatellite loci amplified with the primer pairs nag59F/R, nag111F/R, nag158F/R (**Table S1) (Bell and Ecker, 1994).** We mixed 5-100 ng of template plant DNA in a 50 μL reaction volume, containing 10x Buffer B, 25 mM MgCl2 and 0.5 μL of Taq Polymerase (Biodeal Handelsvertretung Edelmann e.K., Leipzig), as well as 10mM dNTP Mix (Carl Roth GmbH & Co. KG) and 10 μM of each primer of the corresponding primer pair. Products were amplified with the following program: 94°C for 3 min, 40 cycles of 94°C for 15 s, 55°C for 15 s and 72°C for 30 s, followed by a final elongation of 72°C for 3 min. Samples were stored at 4°C until we separated them on 4% Agarose gels (Biodeal Handelsvertretung Edelmann e.K., Leipzig). We identified the size of the fragments via agarose gel electrophoresis and created a phylogenetic tree with this data.

#### Growth rates and flowering times

We sowed *A. thaliana* seeds (NG2, PB, SW1, Woe, JT1 and Col-0) in individual pots on soil (made my mixing 4 L Florador Anzucht soil, 2 L Perligran Premium, 25 g Subtral fertilizer and 2 L tab water) and vernalized them for three days at 4°C in the dark before they were set in the plant growth chamber (PolyKlima, Freising, Germany; growth conditions: 24°C/18°C, 16h/8h day/night cycle at 75% light intensity). For each *A.thaliana* population we randomly chose three of the five plants collected in the wild and used their offspring for the assay. Of each plant three individuals were used, which means that of each plant population we observed nine individuals (n=9). We checked, watered and took pictures of the plants every other day. After 10 days in the plant incubator, we pricked three individual plants of each population and moved them to a new pot with fresh soil. Until the plants grew to senescence we continued to check and water them every other day. We measured the following data: continuous measurements (rosette radius, length of stem), timepoint measurements (flower bud visible for first time, first flower opens, first silique appears) and continuous counting (number of true leaves). Based on these results we assigned growth stages adapted from (Boyes et al., 2001). Seed germination refers to Principal Growth stage 0 and includes seed imbibition, radicle emergence and hypocotyl and cotyledon emergence. Leaf development (Principal Growth stage 2) and Rosette growth (Principal Growth stage 3) occur simultaneously most of the time (more leaves develop while the plants also grow), with rosette growth continuing for a while after the last leaves have developed. The stage Inflorescence emergence (Principal Growth stage 5) begins with the development of a flowering bud and ends when the first flower developed, which is the begin of the Flower development (Principal Growth stage 6). We plotted the data as the mean values of nine individual plant observed of each plant location. The plant population PB needs a second vernalization period for two weeks at 4°C to develop flowering buds. The local population JT1 did not germinate consistently in the lab and was not included in this assay.

#### Leaf glucosinolate analysis

We measured GL profiles of the plants similarly to previously described (Kliebenstein et al., 2001). Four 5-week-old plants of each population were harvested by removing roots and flower stems as close to the rosette as possible. The whole rosettes were transferred into 50 mL tubes, frozen in liquid nitrogen and kept at −80°C until further processing. After freeze-drying for three days, rosettes were grinded to powder by shaking with metal beads for 2 min. GLs were extracted from 8-12 mg of the powder by 1 mL of 80% MeOH with 50 μM pOH-BGl (Sinalbin, internal standard) and thorough vortexing for 30 sec and shaking for 30 min in a paint shaker shaker (Skandex SO-10M, Fluid Management Europe, The Netherlands) at high speed. The samples were centrifuged for 5 min at 3,200 rpm and 600 μL of the supernatant was added to a freshly prepared DEAE-Sephadex-filter plate and allowed to flow through. Samples on the DEAE-Sephadex were washed five times with: (1.) 0.5 mL 80% MeOH, (2./3.) Two times 1 mL ultrapure water, (4.) 0.5 mL 0.02 MES pH 5.2 and (5.) 30 μL sulfatase (Sulfatase from Helix pomatia; Sigma) prepared according to Graser et.al. (Graser et al., 2000). Samples were kept at room temperature overnight and desulfo-GLs were eluted with 0.5 mL dd water on the next day. Desulfo-GLs were then analysed by HPLC-UV. Fifty microliters of the desulfo-GL extract was run on a reversed phase column (Nucleodur Sphinx RP18, 250 x 4.6mm, 5um, Macherey-Nagel, Düren, Germany) on an Agilent 1100 HPLC with a water-acetonitrile gradient (1.5% acetonitrile for 1min, 1.5 to 5% acetonitrile from 1 to 6 min, 5 to 7% acetonitrile from 6 to 8 min, 7 to 21% acetonitrile from 8 to 18 min, 21 to 29% acetonitrile from 18 to 23 min, followed by a washing cycle; flow 1.0 mL min-1). Detection was performed with a photodiode array detector and peaks were integrated at 229 nm. We used the following response factors: 3OHP and 4OHB 2.8, all other aliphatic glucosinolates 2.0, indole glucosinolates 0.5 (Burow et al., 2006) for quantification of individual glucosinolates. The identity of the peaks was based on a comparison of retention time and UV absorption spectrum with data obtained for isolated desulfo glucosinolates as described in Brown et.al (Brown et al., 2003) and by analysis of the desulfoGLs extracts on an LC-ESI-Ion-Trap-mass spectrometer (Esquire6000, Bruker Daltonics). The following glucosinolates were detected in the samples: 3-hydroxypropyl glucosinolate (3OHP), 4-hydroxybutyl glucosinolate (4OHB), 3-methylsulfinylpropyl glucosinolate (3MSOP), 4-methylsulfinylbutyl glucosinolate (4MSOB), 2-propenyl (allyl), 2-hydroxy-3-butenyl glucosinolate (2OH3But), 3-butenyl glucosinolate (3-Butenyl), 4-pentenyl glucosinolate (4-Pentenyl), 4-hydroxy-indol-3-ylmethyl glucosinolate (4OHI3M), 4-methylthiobutyl glucosinolate (4MTB), 7-methylsulfinylheptyl glucosinolate (7MSOH), 8-dmethylsulfinyloctyl glucosinolate (8MSOO), indol-3-ylmethyl glucosinolate (I3M), 4-methoxy-indol-3-ylmethyl glucosinolate (4MOI3M), and 1-methoxy-indol-3-ylmethyl glucosinolate (1MOI3M). Results are given as μmol per g dry weight. Each genotype was tested in four replicates.

To construct the GLs diversity tree, we first filtered out samples with a weight less than 1 mg. The rest of the samples were normalized to the plant weight and combined with the normalized data of a few genotypes from (Kliebenstein et al., 2001) as references. We combined the datasets of both GL analyses and constructed the tree in R using the Neighbor Joining method. In addition, we plot the mean concentrations of each GL.

### Culture-independent characterization of bacterial diversity in wild plant populations

#### Sampling

We sampled the nine populations in February 2019 and 2020, where each sample is made up of 3-4 leaves of one plant, collected with sterilized tweezers (6 replicates per population per time point). Additionally, we sampled six other plants of varying unknown genotypes near each *A. thaliana* plant. In March of 2019 and 2020 we collected *A. thaliana* and other plants from the five selected populations NG2, PB, SW1, JT1 and Woe. We washed all plant samples once with autoclaved ultrapure water to remove dirt and stored them on ice for the remaining sampling time and at −80°C back in the lab. In March 2020 additionally five samples of the soil at the five locations were taken. The upper layer of soil (roughly 1 cm) was removed with a sterilized spatula and then the underlaying soil was scooped in a clean Eppendorf tube. Soil samples were also placed on ice for remaining sampling time and at −80°C back in the lab until further processing.

#### DNA extraction of bacterial communities associated with the wild plant populations

For DNA extraction we transferred the plant material to a 2 mL screw-cap tube with two metal beads (3 mm) and 0.2 g of 0.25-0.5 mm glass beads (Carl Roth GmbH & Co. KG) to crush plant and bacterial cells. Samples were bead-beat two times (30 s at 1,400 rpm on a BioSpec mini bead beater 96) (Biospec Products, Inc, USA) right after they left the freezer to optimize the crushing process. We then added 200 μL CTAB extraction buffer (100 mM TRIS pH 8.0, 20 mM EDTA pH 8.0, 1.5 M NaCl, 2% cetylimethylammoniumbromid, 1% polyvinylpyrrolidone (MW 40,000) in nuclease-free water) to each sample and incubated the samples at 37°C for 10 min. Next, the solution was centrifuged at 20,000 x g at 4°C for 5 min and the supernatants were transferred to a new tube. We precipitated the DNA with 200 μL cold 100% ethanol and centrifuged at 20,000 x g at 4°C for 5 min. We washed the pellet with 70 % ethanol, let it air-dry and resuspended it in 100 μL TRIS-HCl buffer (pH 8). This was followed by a short clean up with home-made Sera-Mag purification beads pre-activated and stored in a PEG/NaCl as described in (Rohland and Reich, 2012).

#### DNA extraction of bacterial communities associated with soil samples at the locations of the wild Arabidopsis thaliana populations

Their DNA was extracted with the “DNeasy Power Soil”-Kit from Quiagen (Qiagen GmbH, Hilden, Germanu). In short, 0.25 g of soil was added to a provided PowerBead tube and vortexed to mix. 600uL of solution C1 was added and the tube was inverted several times. The tubes were vortexed for 10 minutes at maximum speed and then centrifuged for 30 sec at 10,000xg. 400-500 μL of supernatant was transferred to a collection tube. 250 μL of solution C2 was added and samples were vortexed for 5 sec before they were put in the fridge (4°C) for 5 min. Tubes were centrifuged for 1 min at 10,000xg and the supernatant was transferred to a new collection tube. 200 μL of solution C3 was added, samples were vortexed briefly and again incubated for 5 min at 4°C. Tubes were centrifuged for 1 min at 10,000xg and 750uL of the supernatant was transferred to a clean collection tube. 1200 μL of solution C4 was added and samples were vortexed for 5 sec. 675 μL of the whole solution was loaded onto a MB Spin Colum and centrifuged at 10,000g for 1 min. The flow-through was discarded and this step was repeated until the whole samples was processed. 500 μL of solution C5 was added on the column and the sample was centrifuged for 30 sec at 10,000xg. The flow-through was discarded and the sample was centrifuged again for 1 min at 10,000xg. The column was transferred to a clean collection tube and 100 μL of solution C6 was added to the filter membrane. The tubes was centrifuged for 30 sec at 10,000xg and the DNA was used for library preparation.

### Library preparation

We amplified all samples (120 *Arabidopsis* plant samples and 120 “Other” plant samples from 5 locations and 4 sampling time points, 25 soil samples from 5 locations and 1 sampling time point and 3 CTAB buffer controls, 3 negative controls, 3 Zymo (positive) controls) in 10 μL reactions containing 1x Kapa Buffer, 0.3 mM Kapa dNTPs, 0.08 μM of each of the forward primers (341F-OH and 799R-OH; primers with overhang; **Table S1**), 0.25 μM of each of the blocking oligos (Blc_16S_F5 and Blc_16S_R1; **Table S1**) to block amplification of unwanted plant chloroplast DNA, 0.2 μL Kapa taq DNA polymerase (KAPA Biosystems), 4.85 μL nuclease-free water and 2 μL Template DNA (1:4 diluted). The samples were run on a thermocycler block at 95°C for 3 min followed by 5 cycles at 98°C for 20 sec, 55°C for 1 min, 72°C 2 min. The last cycle was followed by a final extension of 72°C for another 2min. We then cleaned the samples enzymatically using Exonuclease I and Antarctic phosphatase (New England Biolabs, Inc.) to degrade leftover primers and inactivate nucleotides (0.5uL each enzyme with 1.22uL Antarctic phosphatase buffer at 37°C for 30 minutes followed by 80°C for 15 min). We used 5uL of the cleaned sample as template for the 2nd (extension) PCR. The 20uL reactions contained as before 1x Kapa Buffer, 0.3 mM Kapa dNTPs and 0.4 μL of Kapa taq DNA polymerase. Each sample received a unique pair of indexed forward and reverse primers in a final concentration of 0.3 μM (Mayer et al., 2021). Samples were run on a similar program as in the first PCR with the difference of the annealing temperature (60°C), a shorter extension time in the cycles (1 min) and for 35 cycles. We cleaned the products with 0.5x vol. of 2x diluted Sera-Mag purification beads and fluorescently quantified the gene products using Picogreen (Thermo Fisher Scientific, Inc.). We used a sample with high fluorescence to prepare a standard curve according to which we calculated the concentrations of the individual samples. We combined the samples in roughly equimolar concentrations and concentrated once more using 0.5x vol. of 2x diluted Sera-Mag purification beads. The final library was quantified with a Qubit (Thermo Fisher Scientific, Inc). The library was denatured and then loaded onto a MiSeq lane spiked with 10% PhiX genomic DNA to ensure high enough sequence diversity. Sequencing was performed for 600 cycles to recover 300 bp of information in the forward and reverse directions with standard Illumina sequencing primers.

#### Processing amplicon sequencing data

We split the amplicon sequencing data on indices and trimmed the adapter sequences from distal read ends using Cutadapt 1.2.1 (Martin, 2011). We then clustered amplicon sequencing data into amplicon sequencing variants “ASVs” using dada2 (Callahan et al., 2016). Reads were first filtered based on their quality (truncLen=c(200,100)) and then dereplicated to eliminate exactly redundant reads. Data was denoised using the inferred error rates ASVs called in the forward and reverse reads. Since not all forward and reverse reads overlapped, we merged forward and reverse reads by keeping overlapping reads and concatenating the rest. We then removed detected chimeric sequences and retrieved a sequence table from the merged data. We assigned taxonomy to the final set of ASVs using the Silva 16S (v 1.32) database (Quast et al., 2013).

### Analysis of microbial diversity

We performed downstream analysis in R with Phyloseq and Vegan (Dixon, 2003; McMurdie and Holmes, 2013). Positive (Zymo microbial community standard) and negative (DNA extraction buffer or no template) controls were as expected (**Figure S2**). Host-derived reads were removed by filtering any ASVs in the order “Chloroplast” and “Ricketsialles” from the 16S ASV tables. Additionally, taxa were agglomerated at genus level. For most analyses (except the bar charts based on relative abundance) we filtered the data to only allow samples with more than 100 reads (without the “Chloroplast” and “Ricketsialles” reads) filtering out 3 samples in total. For barcharts, taxa that were less abundant than 100 times in one sample were removed. For the diversity analysis (beta diversity, beta dispersion and alpha diversity) samples were additionally rarefied to an equal number of reads. Beta diversity is based on the Bray-Curtis distance and ordination performed with principle coordinates analyisis. Statistical significance of explaining variables was tested with an Adonis permutational analysis of variance test. Alpha diversity was inferred using Chao1 and Shannon metrics. The cores of the individual datasets were defined as taxa being present in at least 40% of the samples in each subset. Of the resulting core taxa, the relative abundances were extracted and plotted. Statistical methods used in each analysis are cited in the figures and text.

### Data Availability

All scripts used to generate ASV tables from the raw data, as well as ASV tables, metadata files and scripts that generate the main figures are publicly available on Figshare: http://doi.org/10.6084/m9.figshare.19352189. Raw sequencing data is being made publicly available on NCBI (BioProject ID: PRJNA815825)

## Results

### Whole leaf bacteriomes depend in part on plant species but strong environmental shaping is similar across species boundaries

To study how the assembly of whole leaf bacteriomes are shaped in wild plants, we identified 13 previously unstudied *A. thaliana* populations in Jena, Germany (**Figure S2A and Table S2**). We characterized nine individual *A. thaliana* plants in each location based on length polymorphisms of microsatellite loci (**Figure S2B and S2C**). These polymorphisms suggest that most populations are genetically distinct from one another but have lower diversity within the populations. We sampled plants in nine of these populations by harvesting leaf tissues of both *Arabidopsis thaliana* and other sympatric species (other ground-dwelling ruderal plants randomly selected in the immediate vicinity of the collected *A. thaliana* plants) near the end of February in 2019 and 2020. Each sample is derived from one plant individual. We used ITS amplicon sequencing to characterize the non-*Arabidopsis* plants (Figure S3). Most samples consisted of ITS reads from primarily one plant, resulting in identification to the genus level for most plants. We could not recover ITS reads for some samples and these were not classified. In total there were 25 genera from 12 plant families, and between 4-9 genera per location. The collected species were diverse and included mostly herbaceous plants and grasses that were both annuals and perennials. The exact number of species was not controlled for and varied between sampling events and locations. Therefore, we simply grouped samples for analysis into *A. thaliana* and “other” plants, which resulted in comparable numbers of samples in both groups.

Using 16S rRNA-based amplicon sequencing, we next analyzed the bacterial diversity in the whole leaf samples. Overall, principal coordinates analysis of beta diversity (bacteriome community structures) showed no clean separation of samples based on plant group (*Arabidopsis* vs. other), year or location (Figure 1A). All three factors significantly correlated to beta diversity, but sampling year strongly interacted with sampling location and plant group (Table S3). Indeed, within locations, sampling year was significantly correlated to beta diversity at most sampling sites, indicating strong bacteriome shifts from one year to the next (Table S3). After splitting data into sampling years, we found that the effects of both sampling location and plant host changed from year to year. In 2019, location was significantly correlated to beta diversity (17%, p=0.027) and plant identity was not. In 2020, location was correlated to even more variation (22.9%, p=0.001) and plant identity played a small but significant role (3.1% p=0.003) (Figure 1A).

**Figure 1.**
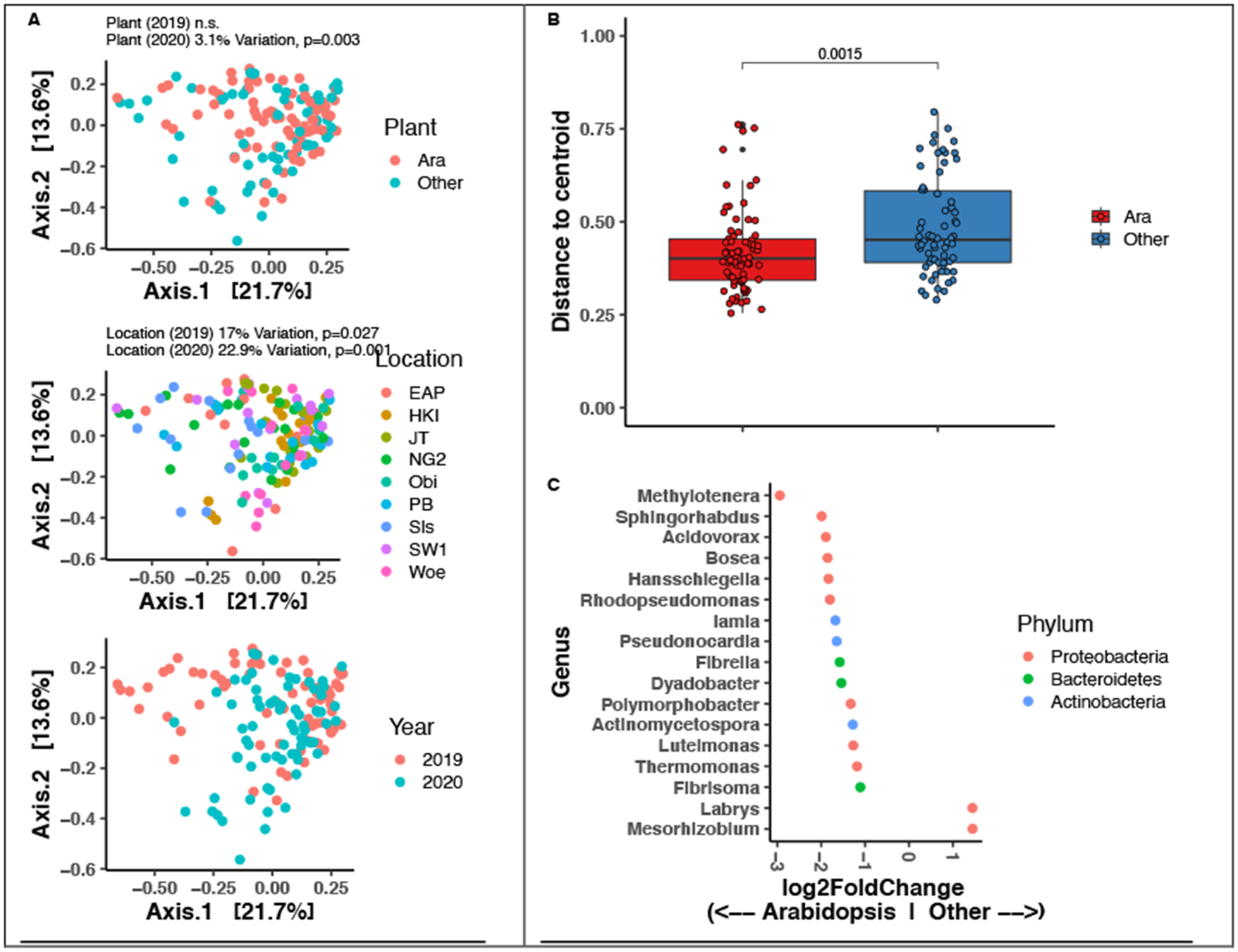
Structure of whole leaf bacteriomes of plants at nine locations in Jena, Germany. **A)** Principle coordinates analysis of Bray-Curtis distances between samples. The three plots are identical but colored by different factors. Percent variation explained by specific factors is shown above the plots (based on a permanova test with 999 iterations). **B)** Plant-to-plant variation in leaf bacteriome community structures, calculated as bray-curtis distance of each leaf sample to the centroid of its group. P-value is based on a t-test. **C)** Bacterial genera that are relatively more abundant in A. thaliana or other plants, determined using a negative binomial model (DESeq2, p<0.001).

We reasoned that comparing *A. thaliana* to mixed non-*A. thaliana* leaf bacteriomes might underestimate the role of plant species. However, if there is a consistent species effect, the leaf bacteriome *A. thaliana* plants should be more similar to one another than the mixed other plant samples are to one another. To test this, we calculated the Bray-Curtis distance between *A. thaliana* samples and between other samples (distance to the group centroids). Indeed, leaf bacteriomes in between individual *A. thaliana* plants were more similar than between other random plants (Figure 1B). This trend was robust within sampling years and within locations (Figure S4). In line with this observation, we were able to detect more taxa enriched in *A. thaliana* than in other plants (Figure 1C). Together, leaf bacteriomes in diverse ground-dwelling ruderals depend on the population location, but the effect of both the location and the plant species varies from year to year.

### A core plant whole-leaf bacteriome common among plant species is clearly differentiated from soil

To better understand the origins and distribution of whole leaf bacterial colonizers, we focused on the most common bacterial colonizers in plants at five selected locations (NG2 - Neugasse2, PB - Paradiesbahnhof, SW1 - Sandweg1, JT1 - Johannistor1, Woe - Wöllnitzer; **Figure 2A).** The *A. thaliana* plants in these populations have high inter-population diversity as measured by microsatellite loci (**Figure S2**). This is supported by both the characteristic profiles of 15 glucosinolates (GLs) in the leaves (**Figure 2B, Figure S5**) and their development times (**Figure 2C**). In these populations the whole leaf bacteriome was characterized in leaf samples collected in both February and March of 2019 and 2020. The bacteriomes were dominated by the phyla *Proteobacteria* (Max. 77% in Woe and min. 63% in SW1), followed by *Bacteroidetes* (Max. 34% in SW1 and min. 12% in Woe; 2020) and *Actinobacteria* (Max. 15% in JT1 and min. 3% in SW1) (**Figure S6, Table S4**).

**Figure 2.**
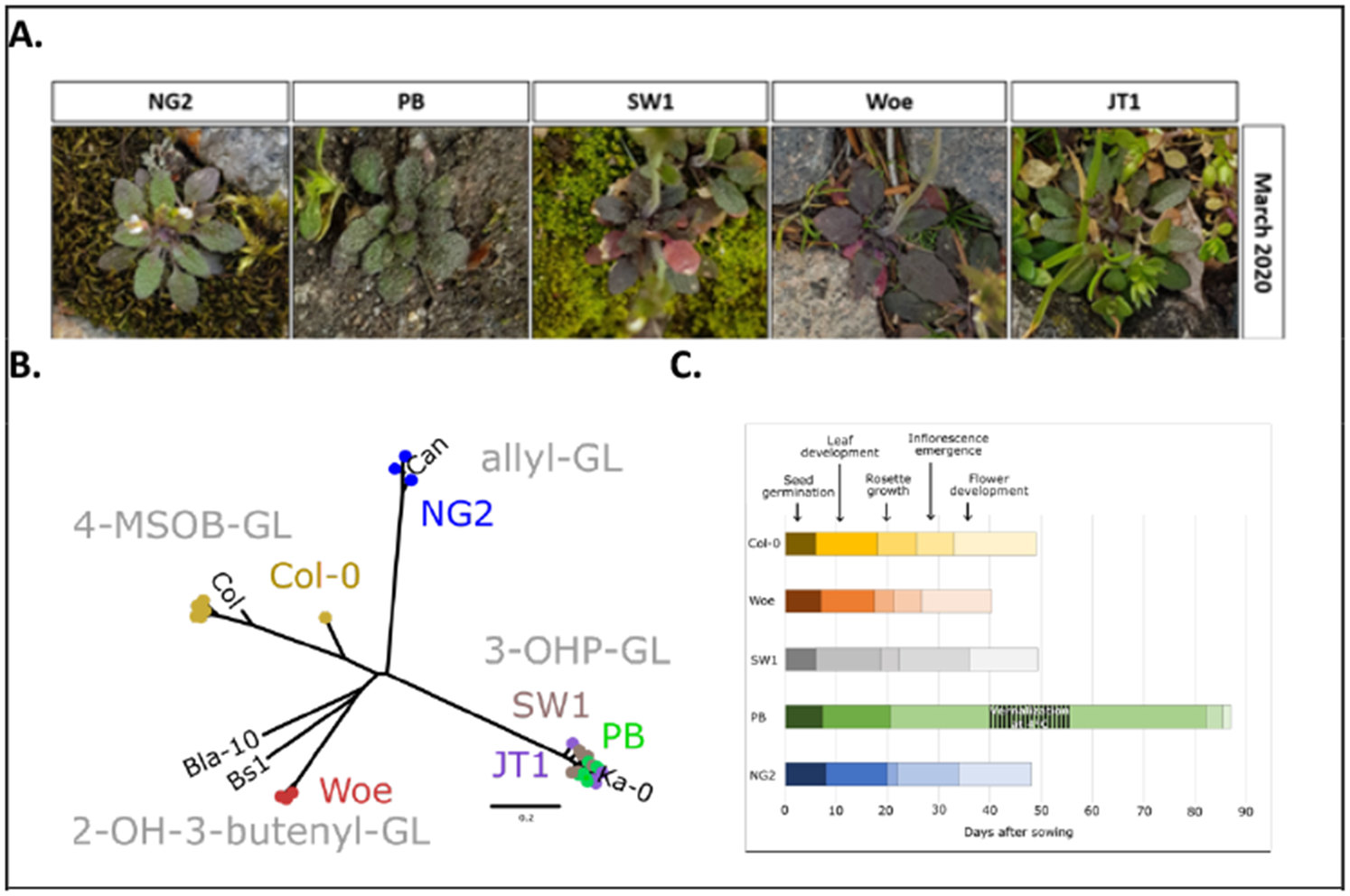
Five diverse A. thaliana populations in Jena, Germany. **A)** Representative *A. thaliana* plants in March 2020. **B)** Diversity of GLs in leaf tissues. The tree is based on Euclidean distance of the normalized GL abundances of four (or three in the case of Woe) repetitions of Jena ecotypes and the reference Col-0 as control. The profiles of reference *A. thaliana* accessions taken from Kliebenstein et al. (Kliebenstein et al., 2001) are shown in black. Each of the five plant populations have one main GL: Col-0 (4-MSOB-GL), NG2 (allyl-GL), Woe (2-OH-3-butenyl-GL) and PB, SW1 and JT1 (3-Butenyl-GL). **C)** Developmental stages of local *A. thaliana* populations. The plant development was assessed based on the growth stages of A. thaliana from (Boyes et al., 2001)) with minor adaptations. Seed germination refers to Principal Growth stage 0. Leaf development (Principal Growth stage 2) and rosette growth (Principal Growth stage 3) occur simultaneously most of the time. Inflorescence emergence refers to Principal Growth stage 5 and Flower development refers to Principal Growth stage 6. The displayed data represents the mean values of nine individual plant observed of each plant location. The plant population PB needs a second vernalization period for two weeks at 4°C to develop flowering buds. Only the population JT1 germinated inconsistently in the lab and was therefore not included in this assay.

To focus on consistent bacterial colonizers, we defined “cores” that appeared in >40% of plant samples, even if they were low relative abundance. We calculated seven cores, one from each location (including both years) and one from each of 2019 or 2020 (including all locations). The overall cores were consistent from 2019 to 2020, with 22 genera in 2019 and 26 in 2020 (**Figure 3A and Table S5**). 19 of the core genera overlapped between the years, including the most prevalent genera *Sphingomonas sp*., *Methylobacterium sp*. *Ellin6055 sp.* and *Hymenobacter sp.* The cores at NG2 consisted of 31 genera, PB 29 genera, JT1 14 genera, SW1 25 genera and Woe 33 genera (**Figure 3B and Table S5**). 11 genera shared at every location were also in the 19 core genera overlapping between years. A further 18 genera overlapped between at least two locations and 21 genera were unique to a particular site. (**Figure 3B** and **Table S5**). Thus, the most common taxa overall were common at all locations and in each year, while a few taxa were more common at certain locations and times.

**Figure 3:**
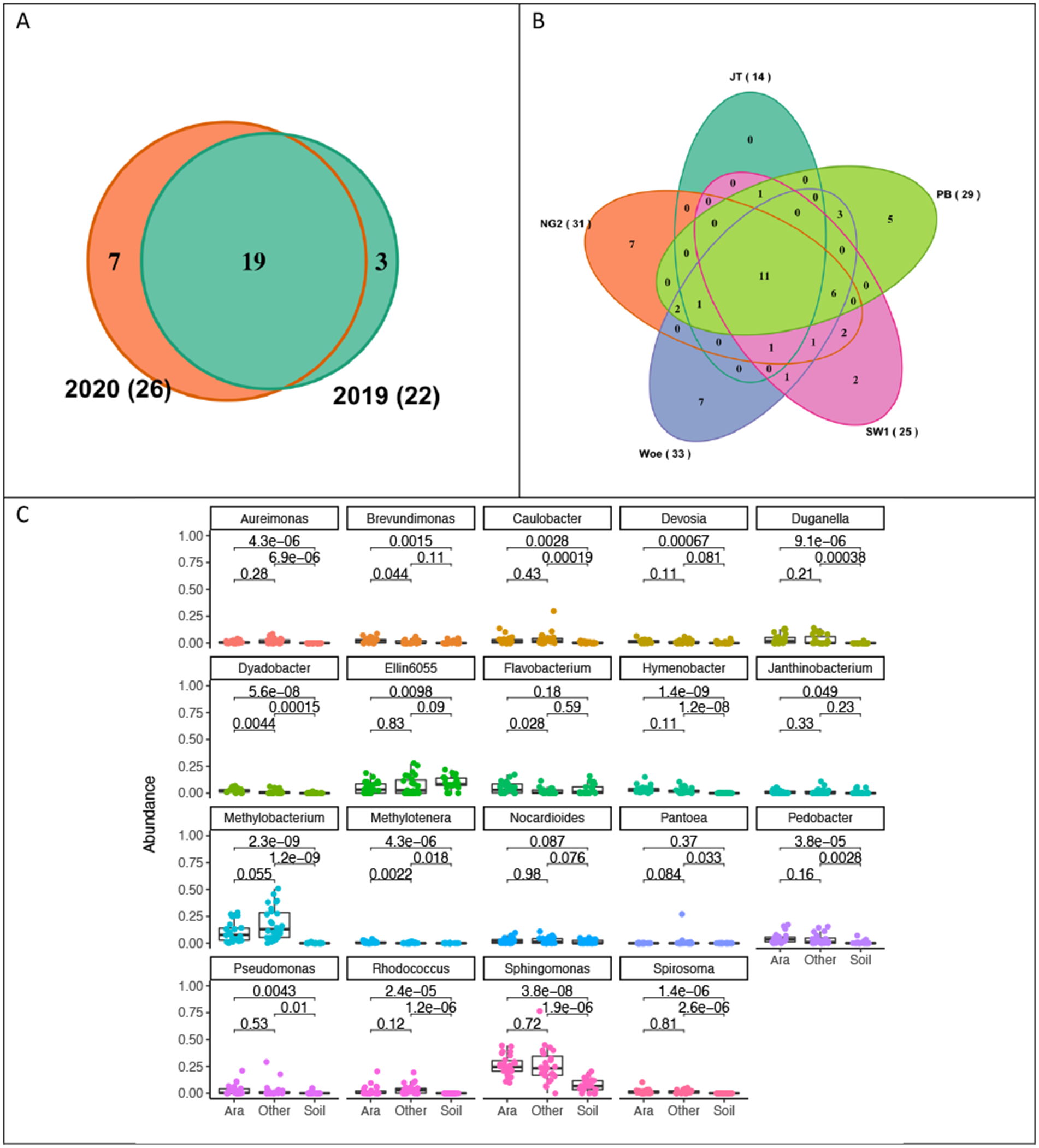
Analysis of the origins and distribution of common leaf bacterial colonizers. **A)** Venn diagram including core microbiomes of two sampling years. In total, the core microbiomes of both years share 19 ASVs. Consequently, there are three ASVs unique to the core from the year 2019 (Bacillus sp., Patulibacter sp., Pantoea sp.) and seven unique to the year 2020 (Janthinobacterium sp., Duganella sp., Ralstonia sp., Nakamurella sp., Mycobacterium sp., Sediminibacterium sp., Bdellovibrio sp.) **B)** Venn diagram including core microbiomes of the five locations. In total, the core microbiomes of the five locations share 11 genera. A further 18 genera overlapped between at least two locations. **C)** Relative abundance of the 19 most prevalent genera of the seven cores calculated in A. thaliana, in other plants and in soil. p-values are based on a Wilcox rank-sum test.

To broadly understand recruitment of these consistent colonizers, we combined the ten most prevalent genera of each year and site core (70 genera total), which considering overlaps resulted in 19 genera. This group thus includes the most prevalent taxa overall (the 11 common to all cores) plus taxa that were prevalent in only some locations or years (all were in at least two cores). We compared the relative abundance of these taxa between *A. thaliana*, other plants and soil in March 2020, when soil samples from the five different locations were collected at the same time as plant samples. Not surprisingly, the soil samples are overall more diverse on phylum level than the plant samples, including seven phyla instead of six (**Figure S7 and Table S6**). There are clear differences between the abundance of the core taxa in the soil compared to the plants. Most (15 of 19) core taxa were detected in soil samples at least once, but many significantly differed in abundance between locations (**Figure 3C and Figure S8**). All of the genera except *Flavobacterium* sp., *Pantoea* sp. and *Ellin6055* sp. were enriched in *A. thaliana* compared to soil. Ellin6055 sp. was most consistently found high abundance in soil samples and was the only taxa significantly depleted in leaves (**Figure 3C**). Interestingly, although non-*A. thaliana* plants had a mix of species, 13 of 19 core taxa were enriched compared to soil at similar abundances as in *A. thaliana*, suggesting overall similar recruitment patterns. Only *Methylobacterium* sp. and *Pantoea* sp. were higher abundance in other plants, while *Brevundimonas* sp., *Dyadobacter* sp., *Flavobacterium* sp. and *Methylotenera* sp. were higher abundance in *A. thaliana* (**Figure 3C**). Together, the most common leaf bacteriome taxa are largely shared among diverse ground-dwelling ruderal plants and are highly enriched compared to the soil they are in contact with.

### Stronger differentiation of whole-leaf bacteriomes in *A. thaliana* populations correlates to lower variability

Next, we looked at just *A. thaliana* leaf bacteriomes at the five local locations to gain deeper insight into formation of location-specific bacteriomes (**Figure 4A**). Overall, leaf bacterial community structure in the *A. thaliana* bacterial communities correlated to location (9.3%, p=0.001), sampling year (5.7%, p=0.001), and sampling month (2%, p=0.019). However, the factors interacted significantly, indicating that location effects are not stable over time (**Table S7**). In all locations, sampling year had a much stronger effect than sampling month, which had little or no effect (**Table S8**). Accordingly, from 2019 to 2020, many taxa became enriched (**Table S9**), corresponding to an increase in alpha diversity (significant in three locations) (**Figure 4B**). Although leaf bacteriomes in 2020 were more diverse, they were also more defined by the sampling location: While in 2019 neither sampling location nor month significantly correlated to bacteriome structure, in 2020 location correlated to almost 20% of beta diversity (p=0.001) (**Figure 4A and Table S10**) with significant differences in alpha diversity (**Table S11**). Similarly, most bacterial genera enriched between locations that could be detected were found in 2020 (**Table S12 and Figure S9**). A more nuanced view of the increase in location specificity can be observed by comparing how similar leaf bacteriomes were to one another within locations vs. between locations. In 2020, samples within locations were more similar than between locations at 4 of 5 locations vs. 2 of 5 in 2019 (based on Bray-Curtis distances, **Figure 4C**). This was caused by a major decrease in variability in 2020 at locations JT1, NG2 and PB (**Figure S11**, all p < 5×10^−5^) and an increase only at SW1 (**Figure S11**, p=0.008). Thus, while leaf bacteria at most locations were more specific in 2020, this was not universal.

**Figure 4:**
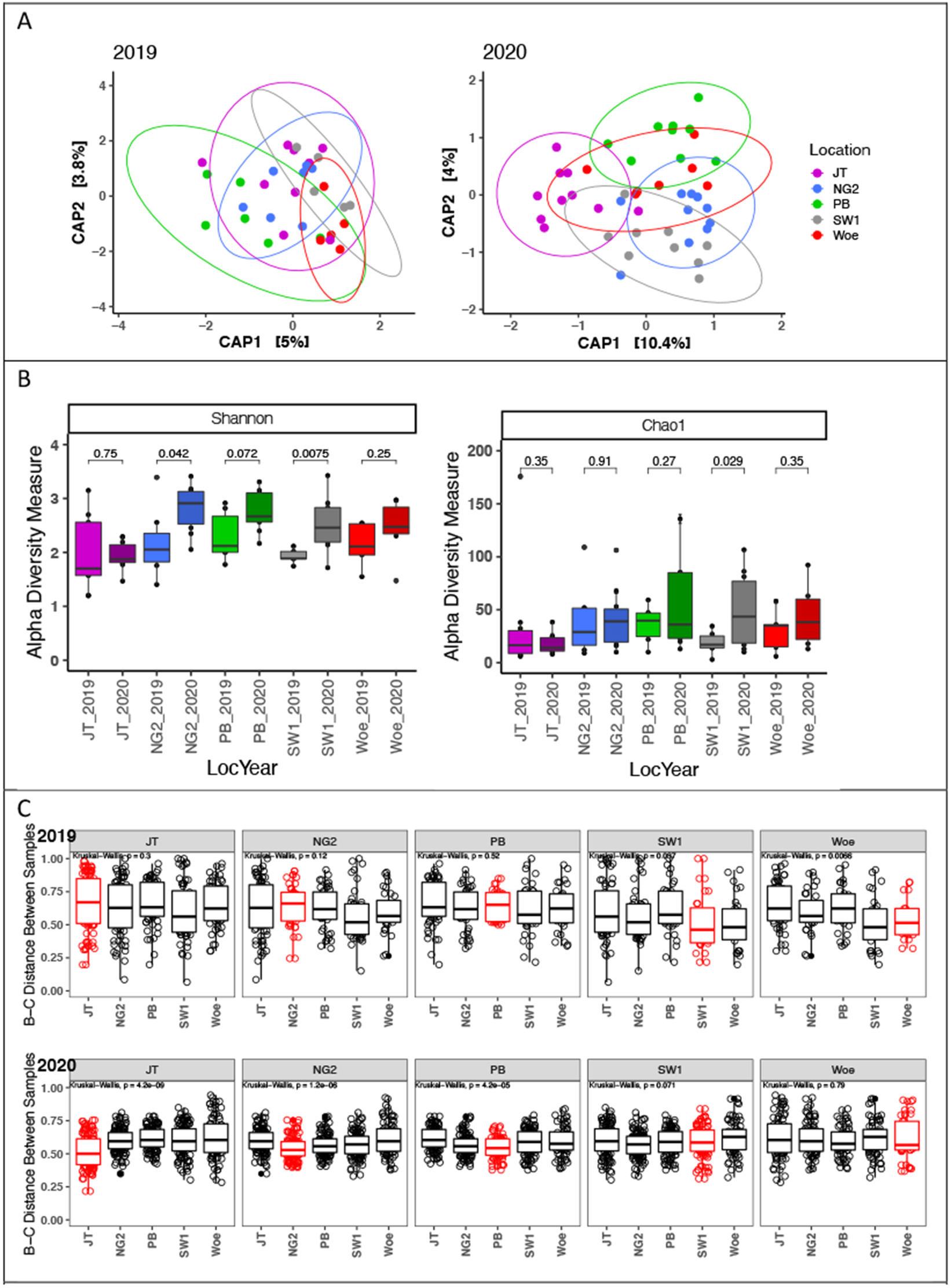
Diversity of leaf bacteriomes in A. thaliana plants collected in Feb and March 2019 and 2020. **A)** Ordination of bray-curtis distances between samples, constrained by location in 2019 and 2020. Betadiversity differed significantly between the locations in 2020 and contributed to 20% of the observed variation. It did not differ significantly in 2019. (Permanova) **B)** Alpha diversity (Shannon and Chao1) of leaf bacterial communities by location and year (p-values based on t-test). Alpha diversity tended to increase in 2020 (significant at 3 locations) and differed between locations only in 2020 (see also Table S11) **C)** Bray-curtis distances between samples at the same location (red) vs between samples from different locations (black). Each plot compares samples of one location (plot header) to samples of the same or other locations (x-axes). Samples were overall generally more similar within locations in 2020.

Importantly, the trends in plant-to-plant variation and leaf alpha diversity observed in 2020 were distinct from local soils: While soil bacterial communities at different locations were significantly different (explaining 25.7% of variation, permanova p=0.003), in contrast to leaves, soil alpha diversity did not significantly differ between locations and distance between samples was lowest in NG2 and highest in JT1 (**Figure S12 and Table S13**).

Taken together, leaf bacteriomes in *A. thaliana* populations are more location- and host-specific (less variable within groups) in some years than others (see also **Figure 1**) with the direction of observed trends depending on the population.

### Leaf chemotype strongly correlates to level of leaf bacteriome variability

Since locations (environments) are in this case fully nested with *A. thaliana* ecotypes, we cannot distinguish whether different patterns in leaf bacteriome recruitment could depend on host genotypes. Therefore, we turned to leaf traits. At seven locations where we characterized *A. thaliana* whole leaf bacteriomes, we also measured indole and aliphatic GL profiles (**Figure S5**). While indole GLs were similar across ecotypes, the main aliphatic GL profiles were distinctly different. In Europe, seven different aliphatic glucosinolate chemotypes are known whose composition is determined by only three genetic loci (Katz et al., 2021). Plants of different aliphatic chemotypes usually have one main aliphatic GL with very little overlap of individual compounds, which was also true for our plant populations (**Figure S5**). We observed three of seven chemotypes: allyl-GL at NG2, 3-OH-Propyl-GL (hereafter 3OHP) at PB, SW1 and JT1 and 2-OH-3-butenyl-GL (hereafter 2OH3But) in Woe, SLS and EAP. Allyl-GL and 3OHP-GL are common in Germany, whereas 2OH3But-GL is more common in other parts of Europe (Katz et al., 2021). Since aliphatic glucosinolates have been shown to play important roles in bacterial colonization (Fan et al., 2011), we checked whether we could detect effects of leaf chemistry independent of sampling locations using the three 3OHP and three 2OH3But populations (all sampled in February 2019 and 2020).

As expected, whole leaf bacterial community structures overall significantly correlated to year (6.8%, p=0.001), with significant interactions between year and chemotype (5%, p=0.002). Therefore, we divided the data into years and found that chemotype only significantly correlated to bacterial community structures in 2020 (**Figure 5A and 5B**, 9.3%, p=0.003). A DESeq2 differential abundance analysis suggested that in 2019, 12 genera were significantly enriched (p<0.05) between chemotypes and in 2020 that one genus was enriched. Of those, only *Bacillus* sp. in 2019 and *Sphingopyxis* sp. in 2020 (both enriched in 2OH3But), were significantly enriched when evaluated using a stringent Wilcoxon rank-sum test (Figure S13).

**Figure 5.**
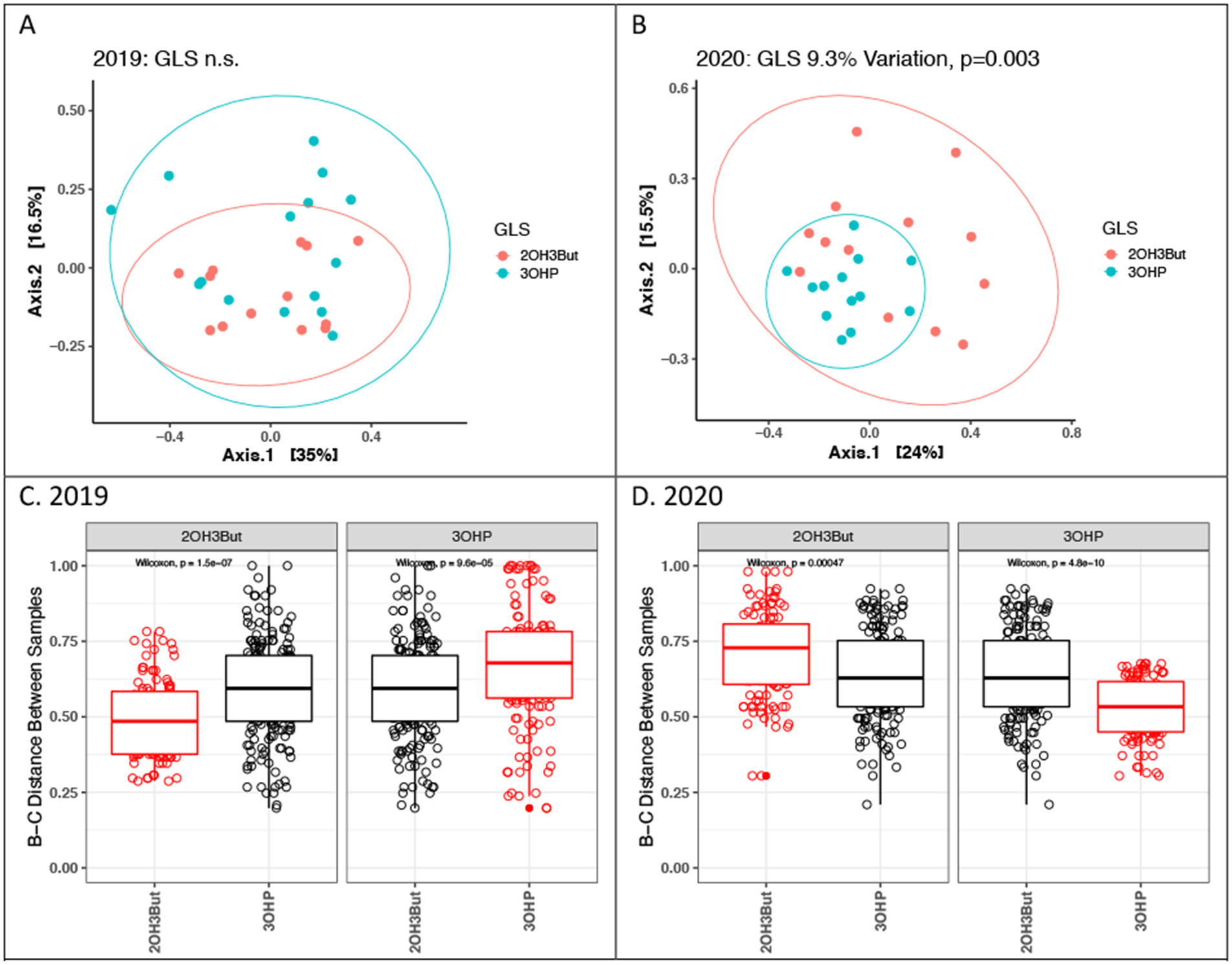
Beta diversity analysis of leaf samples from three locations with 2OH3But and three locations with 3OHP chemotypes. **A and B)** Principal coordinates analysis of bray-curtis distances between samples suggests a significant effect of chemotype only in 2020. Percent variation explained and p-values calculated with a permanova analysis with 999 permutations. Ellipses represent 95% confidence intervals. **C and D)** Bray-curtis distances of leaf bacteriomes between samples sharing a chemotype (red) vs between samples with different chemotypes (black). Comparisons between samples from the same location are excluded. Each plot compares samples of one chemotype (plot header) to samples of the same or other chemotype (x-axes). Effects of chemotype changed between 2019 and 2020. P-values are based on a Wilcoxon rank-sum test.

The ordinations did appear to show that the bacteriomes of one or the other chemotype were more variable in each year. Thus, we measured the similarity of bacterial communities in leaves sharing a chemotype and in leaves with different chemotypes. Since location has strong effects, to make a fair comparison, we excluded comparisons between samples at the same location. Indeed, in 2019, 2OH3But leaf bacteriomes were significantly more similar to one another than they were to 3OHP leaf bacteriomes (p= 1.5×10^−7^), while in 2020 this was true of 3OHP bacteriomes (p=7.3×10^−7^) (**Figure 5C and 5D**). To our surprise, in each year, leaf bacteriomes in the other chemotype (3OHP in 2019 and 2OH3But in 2020) were significantly *less* similar to one another than to plants in populations with a different chemotype (p=9.8×10^−5^ and 0.00045, respectively) (**Figure 5C and 5D**). Therefore, our data suggests a possible role for leaf chemotypes in constraining leaf community structures *and/or* in promoting beta diversity of leaf community structures. Importantly, the outcome is apparently dependent on an unknown factor that changed between growing seasons.

## Discussion

Microbial communities associated with the leaves of plants are diverse and composed of bacteria, fungi, oomycetes and other eukaryotes (Mayer et al., 2021) that play key roles in host health (Liu et al., 2020). Therefore, understanding host controls on leaf colonization processes and the role the microbiome plays in the local adaption of plants will be important in developing strategies to shape plant microbiomes in beneficial ways. One approach to do this is to use panels of diverse plants in garden experiments, which can provide valuable insights into colonization under controlled conditions (Horton et al., 2014) or to use reciprocal planting experiments to gain insights under partially controlled conditions (Wagner et al., 2016). While powerful, such experiments likely cannot fully capture natural dynamics since wild plants over time build up required associations in response to local context, including disease threats and polyculture with other plants (Santhanam et al., 2015). Therefore, wild plant populations can still serve as an important tool for studying the assembly of plant microbiomes (Agler et al., 2016). To address the challenge of understanding how host factors shape microbiomes, *Arabidopsis thaliana* is a valuable tool due to persistent wild populations, usually co-colonized by other plants, and diverse physical and chemical phenotypes, sometimes across small geographic scales (Katz et al., 2021; Seren et al., 2017; Weigel, 2012).

Soil is a primary source of inoculum for both roots and leaves of plants (Massoni et al., 2021; Tkacz et al., 2020) and probably continuously inoculates ground-dwelling ruderal plants due to their contact with the ground and splashing during rain events. Soil bacteria are extremely diverse (Ramirez et al., 2014) but we observed similar to previous reports that leaves of *A. thaliana* tend to be colonized mainly by bacteria belonging to a few phyla, including *Proteobacteria*, *Bacteroides* and *Actinobacteria* (Bodenhausen et al., 2013) Consistent with previous work (Wagner et al., 2016), most bacteria that consistently colonized leaves were also observed in soil samples but in differential abundance. Many taxa were extremely enriched compared to soil, such as *Sphingomonas* sp., *Hymenobacter* sp. and *Rhodococcus* sp., underscoring strong plant selection and their fitness as leaf colonizers. Other taxa like the common soil taxa *Ellin6055* sp., while still common in leaves, was selected against in the leaf bacteriome. Interestingly, we found that all common *A. thaliana* colonizers were also found on other plants, in most cases at similar abundances. Previous observations similarly showed in three herbaceous plant species that 98% of identifiable taxa were shared with similar prevalence across hosts (Massoni et al., 2020). Thus, most wild ground-dwelling ruderal plants in these populations appear to have conserved colonization dynamics leading to clearly distinguished bacteriomes from the soil but similar bacteriomes in different plant species and one location.

We did observe some differences between hosts, in *Flavobacterium*, *Brevundimonas* sp., *Methylobacterium* sp. and *Methylotenera* sp., the latter two of which are facultative methanotrophs that can grow on C1 compounds (Kalyuzhnaya et al., 2012). Together with previous results observing similarly that *Methylobacterium* sp. community composition is significantly affected by plant host species (Knief et al., 2010), C1 metabolism seems to be an important factor underlying differentiation of multiple prevalent leaf colonizers. This could be important, since at least *Methylobacterium* sp. can provide plant-beneficial effects (Grossi et al., 2020). *Flavobacterium* sp. was previously shown in lab colonization experiments to be differentially abundant between plant species rhizosphere microbiomes (Wippel et al., 2021), but the reasons for their differentiation in leaves will need more research. Recently, the same strains of *Pseudomonas* sp. and *Sphingomonas* sp. were shown to thrive in the wild in leaves of both *A. thaliana* and other plant hosts (Lundberg et al., 2021). Such approaches that can distinguish bacteria at finer levels are needed because they can lead to better understanding of plant recruitment strategies in nature.

Besides effect of host plant species, we observed a strong effect of sampling location. Differences between populations in *A. thaliana* have been observed before (Agler et al., 2016) and many factors could contribute. For example, NG2, PB and JT1 are near the city center of Jena, and an urban/rural divide has previously been suggested to influence tree-leaf bacterial communities (Laforest-Lapointe et al., 2017). Whatever the cause of inter-location effects, they were inconsistent: we observed an increased effect of location from 2019 to 2020 in both *A. thaliana* (~11% to ~20%) and in all plants (from 17% to 23% of variation). Similarly, the effect of both host plant species and leaf glucosinolate chemistry on leaf bacterial communities also increased in 2020. A shift in 2020 could also be observed within locations where sampling year correlated to 11.7% to 15.7% of observed variation. Importantly, we observed not just a shift in strength of explanatory variables, but also that this was due to changes in plant-to-plant bacteriome variation. Phyllosphere microbiota are known to have temporal dynamics, most likely due to environmental conditions and nutrient availability (Copeland et al., 2015). The winter of 2019-2020 (the growing season of *A. thaliana*) was abnormally warm and dry with no days below freezing and frequent temperature spikes up to 15oC (data for the Jena Sternwarte weather station, DWD climate data center, https://opendata.dwd.de/climate_environment/CDC/). Thus, we hypothesize that climate could be to blame for the effects in 2020.

Temperature could affect leaf bacteriome assembly in multiple ways. In our ecotypes, development times under lab conditions were highly divergent. In an unusually warm winter, development could become magnified because of missing temperature cues. The absence of winter temperature spikes was previously shown to be important in regulation of flowering in *A. thaliana* (Hepworth et al., 2018) and could regulate other processes like temperature-dependent phyllosphere microbial community assembly (Aydogan et al., 2020). Additionally, since we sampled individual plants, we could observe that the shift in 2020 to more differentiated leaf bacterial communities was caused by a strong decrease in the variation in leaf bacteriomes at several locations and an increase only in one location. Our results suggest that this population-specific behavior might in part be explained by leaf chemotypes. If so, it is reasonable to expect temperature to play a role, since production of glucosinolates (GLs) themselves are regulated by temperature in *A. thaliana* (Kissen et al., 2016). Additionally, upon leaf damage GLs encounter myrosinase enzymes and break down to a range of potentially toxic products (Halkier and Gershenzon, 2006). For example, sulforaphane, the isothiocyanate product of 4-MSOB GLs, regulates bacterial colonization by inhibiting type three secretion system expression (Wang et al., 2020). The exact hydrolysis products of each chemotype are different and further depend on temperature (Hanschen et al., 2017), which likely comes with differences in volatility and biological activity. Thus, GLs could reasonably account for temperature-dependent bacteriome differences.

An open question then is would plant ecotypes regulate plant-to-plant variation in leaf microbiomes or are colonization phenotypes just a passive effect of environment and for example plant chemistry? Plants in the wild must deal with unpredictable threats and therefore, phenotypic variation is critical survival and adaptability. Leaf microorganisms play important roles in defining plant traits and thus can extend plant phenotoypes (Hawkes et al., 2021). Therefore, increased plant-to-plant leaf microbiome variation could increase chances of survival. This would not come without risk, since essentially allowing or even driving random colonization can be detrimental. Thus, a regulated system that auto-tunes according to environment might be a good strategy to balance adaptability and risks. At any rate, our results show that factors shaping leaf bacteriomes exert effects that probably depend on environmental conditions, but more work is needed to understand whether and how this affects plant health. Gaining this understanding, however, could help design better approaches to harness microorganisms for sustainability and should be an important focus of ongoing research.

## Acknowledgements

TM and MTA were supported by the Carl Zeiss Stiftung via the Jena School for Microbial Communication and the Friedrich Schiller University of Jena. They were also funded by the Deutsche Forschungsgemeinschaft (DFG, German Research Foundation) under Germany’s Excellence Strategy – EXC 2051 – Project-ID 390713860. MR and JG were supported by the Max Planck Society.

## Author Contributions

MTA conceptualized the project. TM performed plant sampling and other analyses and generated sequencing data. MR analyzed the glucosinolate composition of leaf samples. TM designed and wrote the scripts to process data and generate figures with help from MTA. TM and MTA wrote the manuscript and all authors edited and approved it.

