## Supplementary Information for "Leaf bacterial community structure and variation in wild ruderal plants are shaped by the interaction of host species and defense chemistry with environment"

Supplementary Figures S1 – S13

Supplementary Tables S1 – S13

Supplementary Files:

File S1 & R-script 1 – Data and Script to create Microsatellite Tree

File S2: Excel File with data of Flowering Time assay

File S3 & R-script 2 – Data and Script to create Glucosinolate Tree

File S4: Sequences of the barcoded primers used in this study (library preparation)

File S5: Core ASVs of the years and the locations

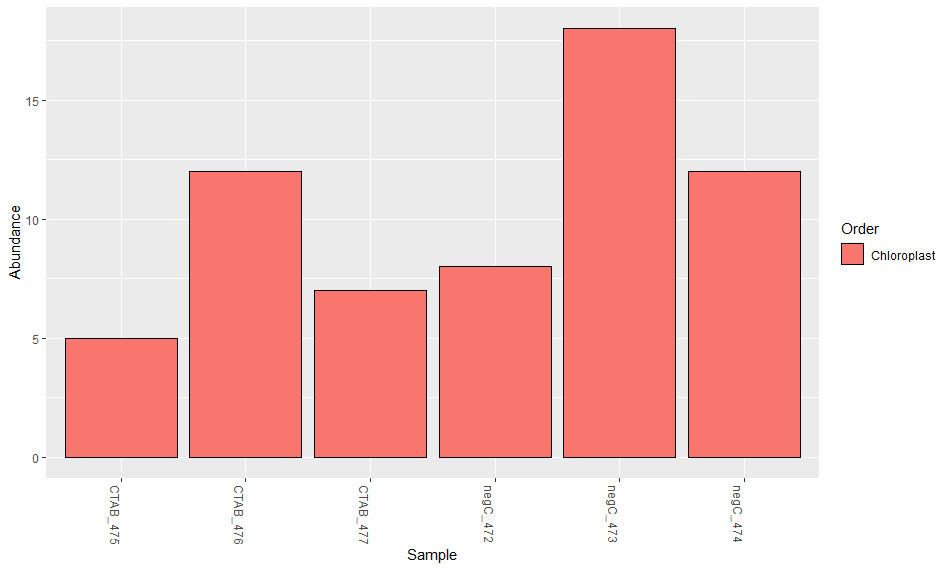
A)

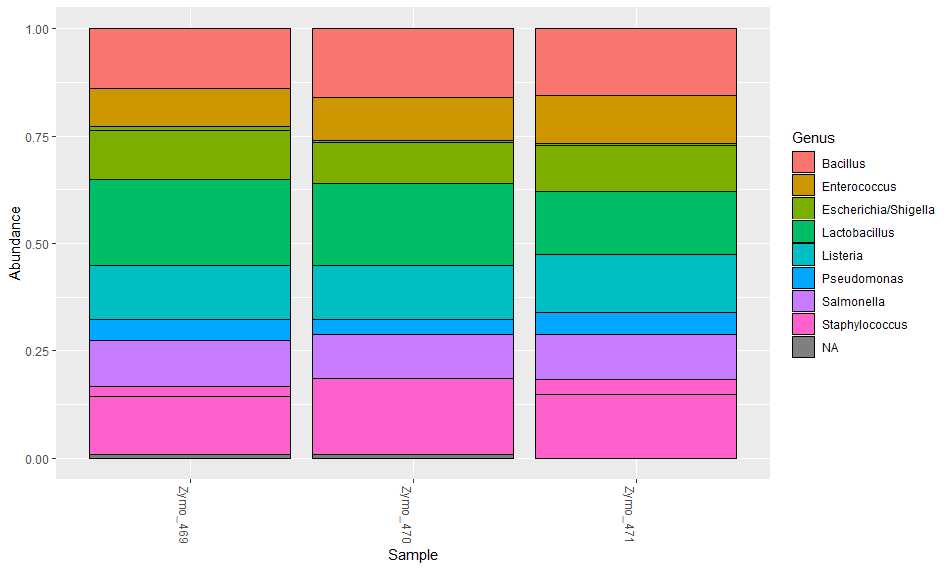
B)

**Figure S1: Control samples. A) CTAB and negative control.** Three replicates of the CTAB buffer used for bacterial DNA extraction from plant leaves as well as three replicates of nuclease-free water as PCR control were sequenced in addition to the other samples. As expected they yielded very low read counts (from 5 to 16 reads per samples). **B) Zymo control.** As positive control served the ZymoBIOMICS microbial community DNA standard (Zymo Research) containing a mix of 10 microbes including 8 bacteria and 2 fungi in . All 8 bacteria were detected, however not in the equal abundances as they are supposed to be in the Zymo Mix.

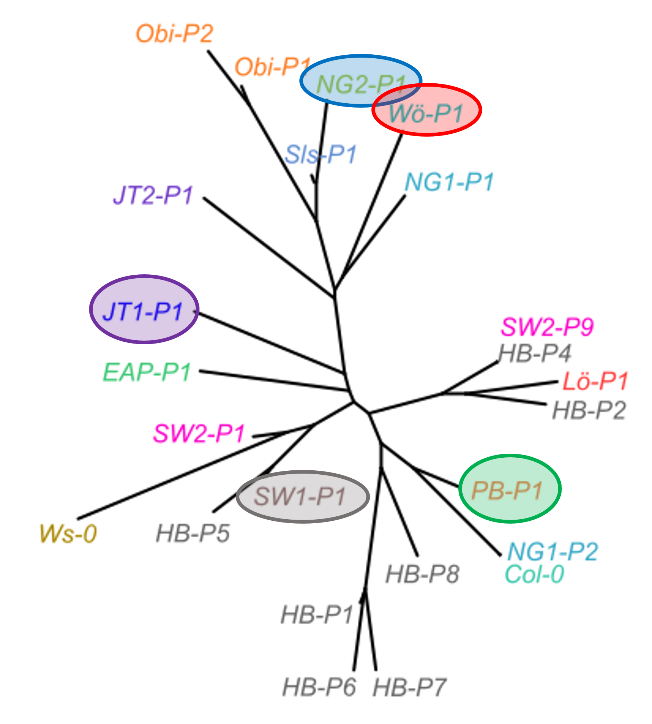

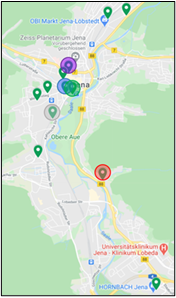
**A. C.**

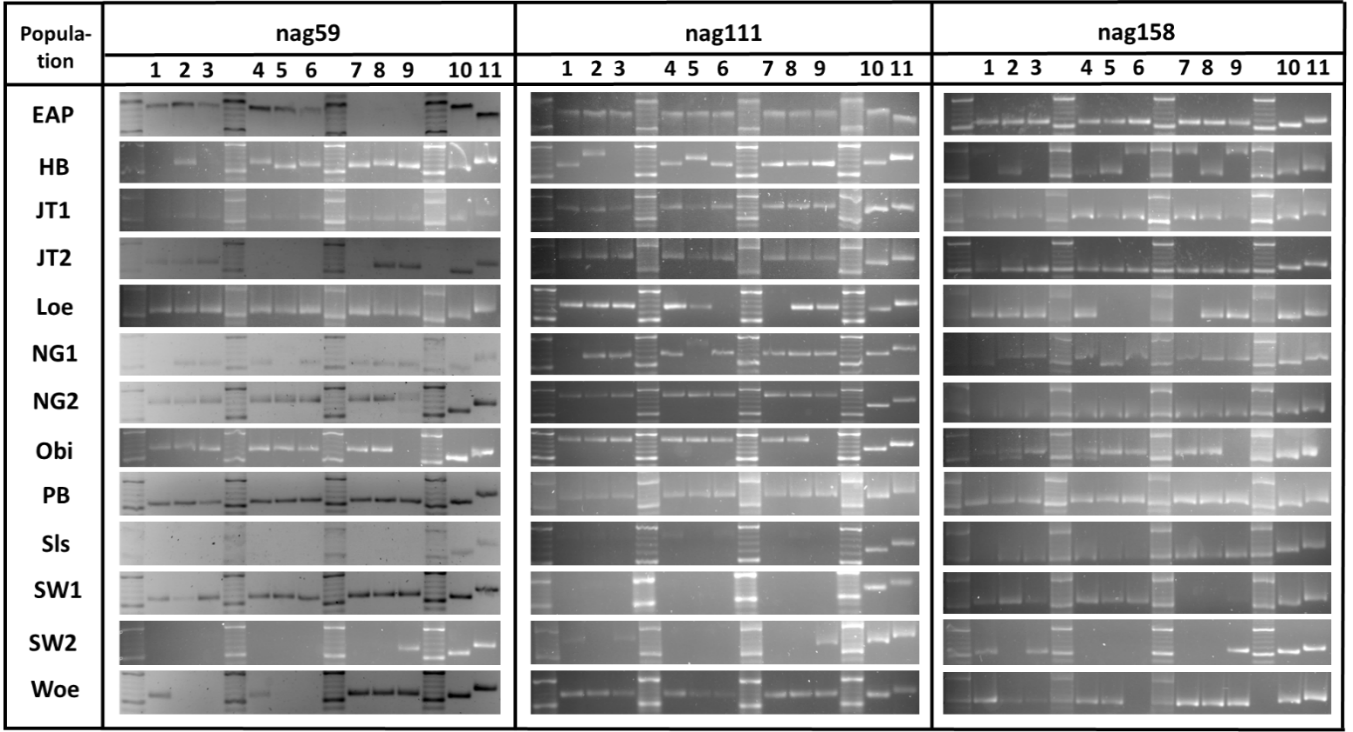
**B.**

**Figure S2: Local A. thaliana populations in Jena, Germany.** (A.) In spring 2018, we identified 13 local A. thaliana populations (see also **Table S1**). (B.) Length polymorphisms in three microsatellite loci after PCR amplification to observe plant diversity. The gel-cutout shows the region between 200 bp (upper thicker band of the ladder) and 100 bp (lower thicker band of the ladder). Col-0 (lane 10) and Ws-0 (lane 11) were added as a control to each PCR. The populations NG2, PB, SW1, Woe, JT1, EAP, Sls and Loe showed uniform microsatellite profiles in all plants. The populations NG1, SW2, JT2, Obi and HB had at least one plant with a varying microsatellite profile compared to most of the populations profile (C.) The microsatellite data was used to calculate a neighbor-joining tree (see supplementary methods). Five populations (NG2, PB, SW1, Woe, JT1) were widespread on the microsatellite tree (high inter-population diversity) but had uniform microsatellite profiles (low/no intra-population diversity).

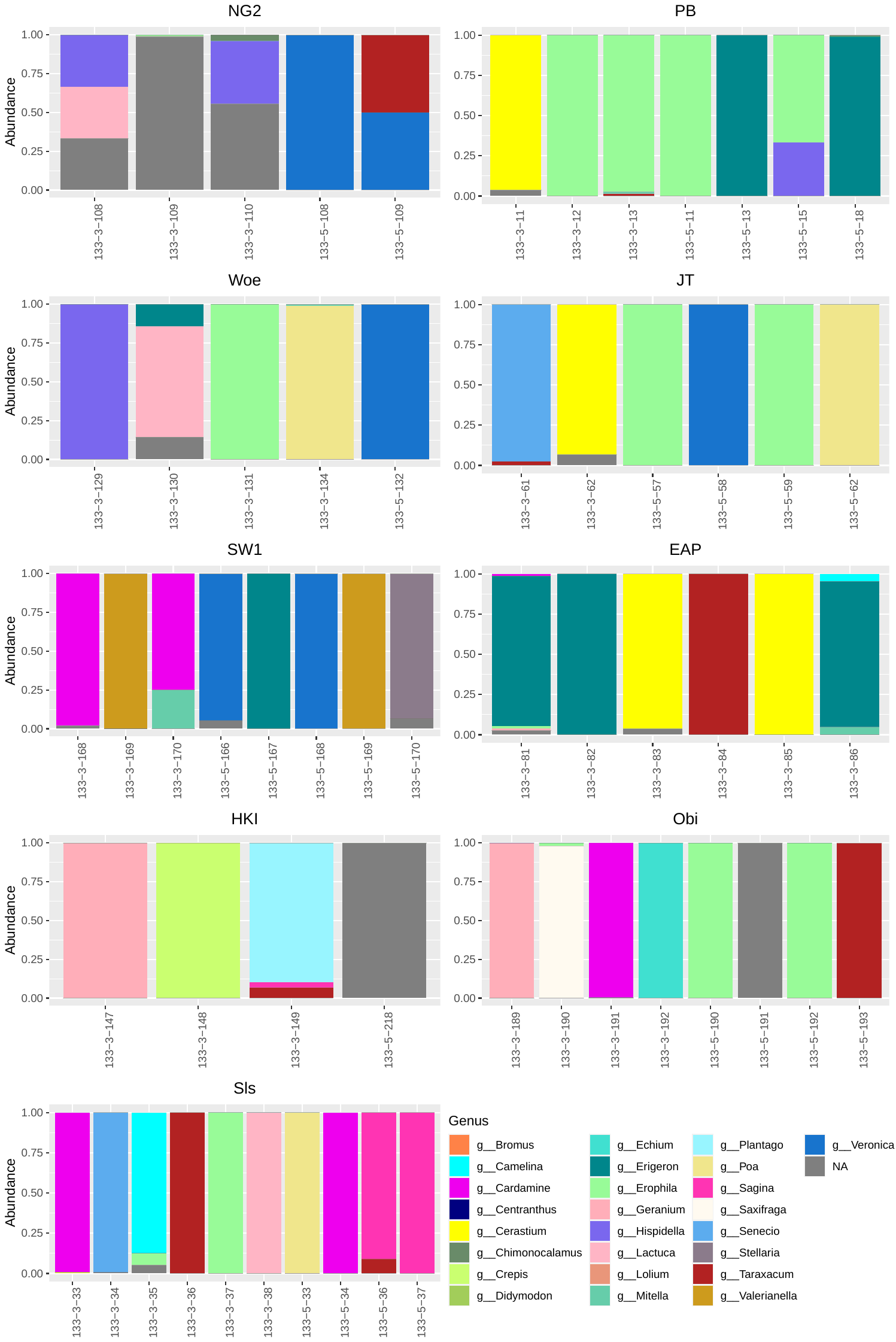

**Figure S3:** Identification of “other” plant samples collected from the sites of nine *A. thaliana* populations at the genus level using ITS sequencing data. Samples are only shown for which ITS data were recovered, which does not reflect all samples collected and used for 16S rRNA sequencing.

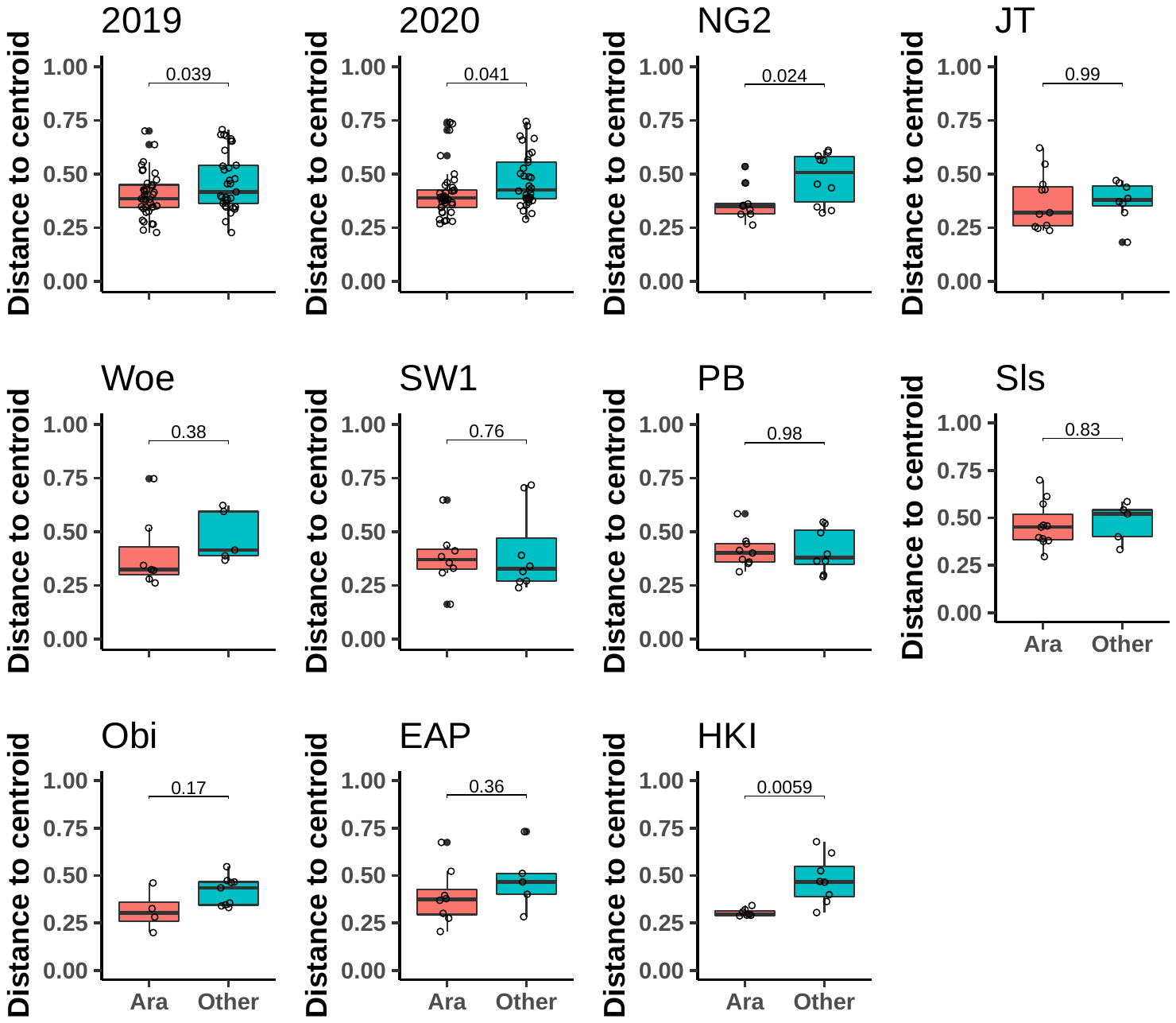

**Figure S4:** Comparison of bray-curtis distances between whole leaf bacteriomes in *A. thaliana* plants and in other plants. Distance of samples to the group centroid, calculated within years or within locations. All significant differences and most trends show that *A. thaliana* leaf bacteriomes are more similar to one another than those of other plants.

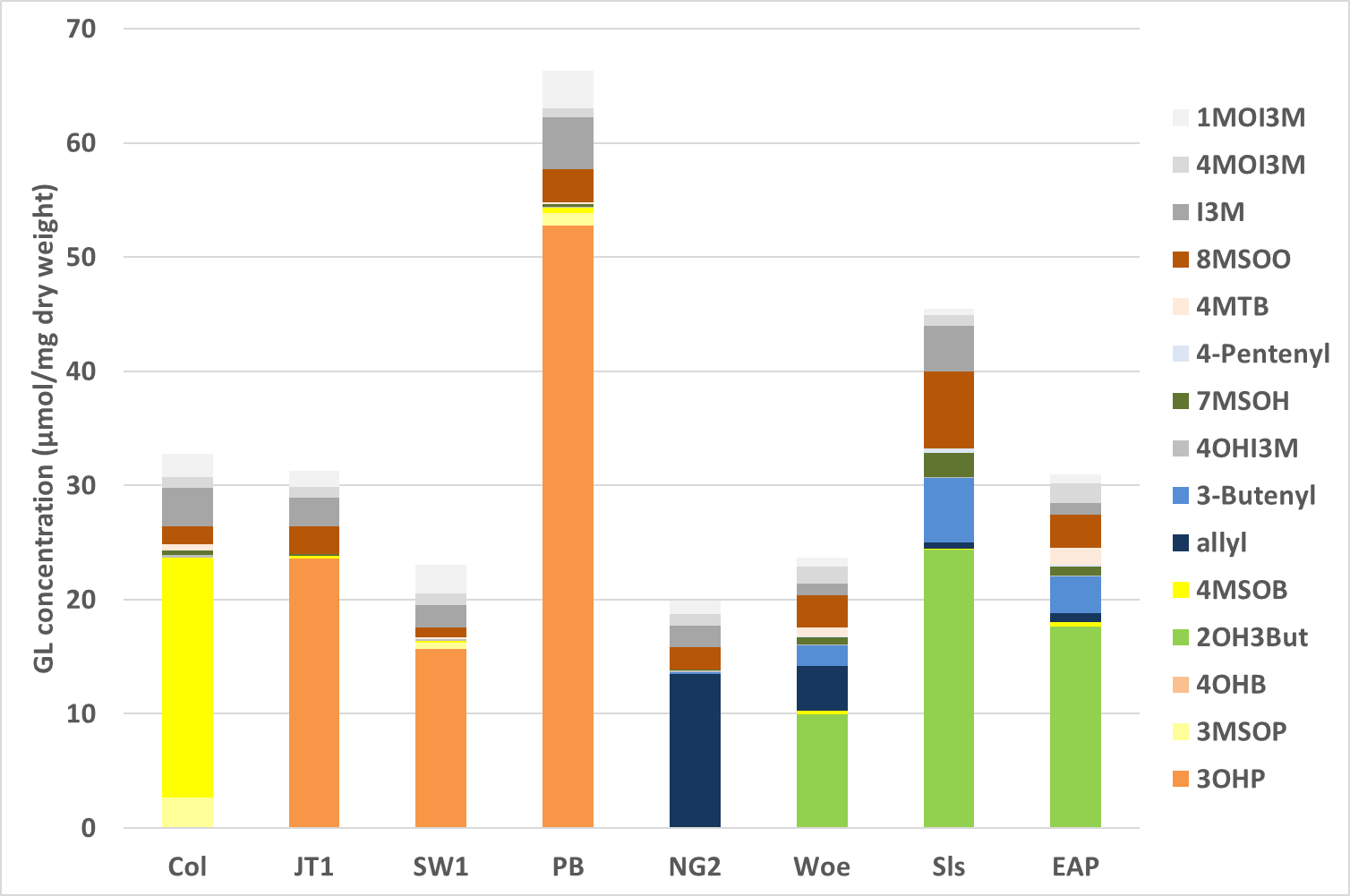

***Figure S5: Concentrations of glucosinolates in leaves of plants from seven different populations.*** *The bar chart shows the aliphatic GLs in color and the indole GLs in gray shades. Each plant population has one main GL, which are different NG2 (allyl-GL), Woe, SLS and EAP (2-OH-3-butenyl-GL) and PB, SW1 and JT1 (3-hydroxypropyl-GL). Absolute amounts also differ: Total GLs levels were highest in PB and lowest in NG2. Col-0 (4MSOB-GL) is a commonly used reference genotype measured as control.*

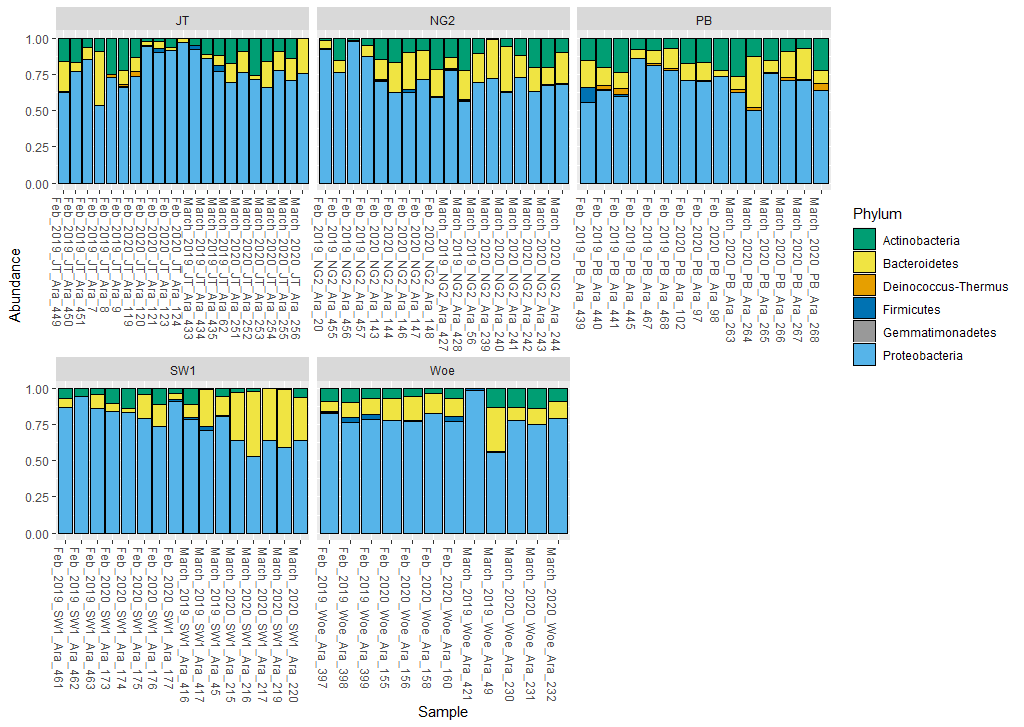

**Figure S6: Identities of the bacterial communtities of the five locations in the two different years on phylum level**. Taxa that were presented less than 100 times in the dataset were removed. Displayed are the barplot for each individual sample of the different years, months and locations. The most heavily sequenced phylum was Proteobacteria, followed by Bacterioidetes and then Actinobacteria. The phyla Deinococcus, Firmicutes and Gemmatimonadetes were only represented 0-2% in the datasets.

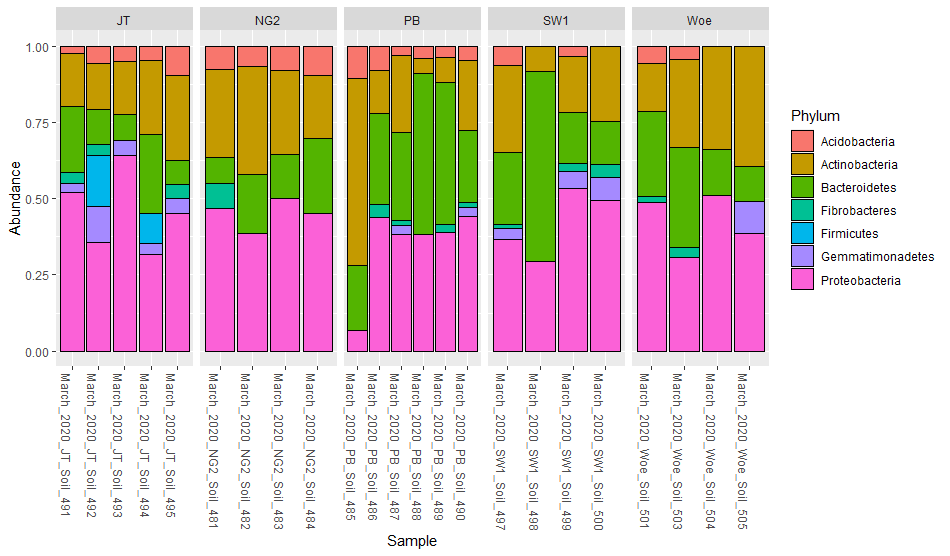

***Figure S7: Bacterial communities of the soil samples of the five locations on the phylum level.*** *Taxa that were present less than 100 times in the dataset of one year were removed. Displayed is the barplot for each individual sample. The most heavily sequenced phylum was Proteobacteria, followed by Bacterioidetes and then Actinobacteria. Abundances of Acidobacteria and Gemmatimonadetes range from 0 to 6%, and abundances of Fibrobacteria from 2-3%. Firmicutes are only found in JT1 (7%). Like in the plant samples, Proteobacteria is the most abundant phylum, however it only reached a maximum of 46% compared to maximum 86% in leaf samples. The second most abundant phylum is Bacteriodetes (Max: 36% in PB, min.: 16% in JT1), followed by Actinobacteria (Max.: 29% in Woe, min.: 19% in PB). Abundances of Acidobacteria and Gemmatimonadetes range from 0 to 6%, and abundances of Fibrobacteria from 2-3%. Firmicutes were only found in JT1 (7%).*

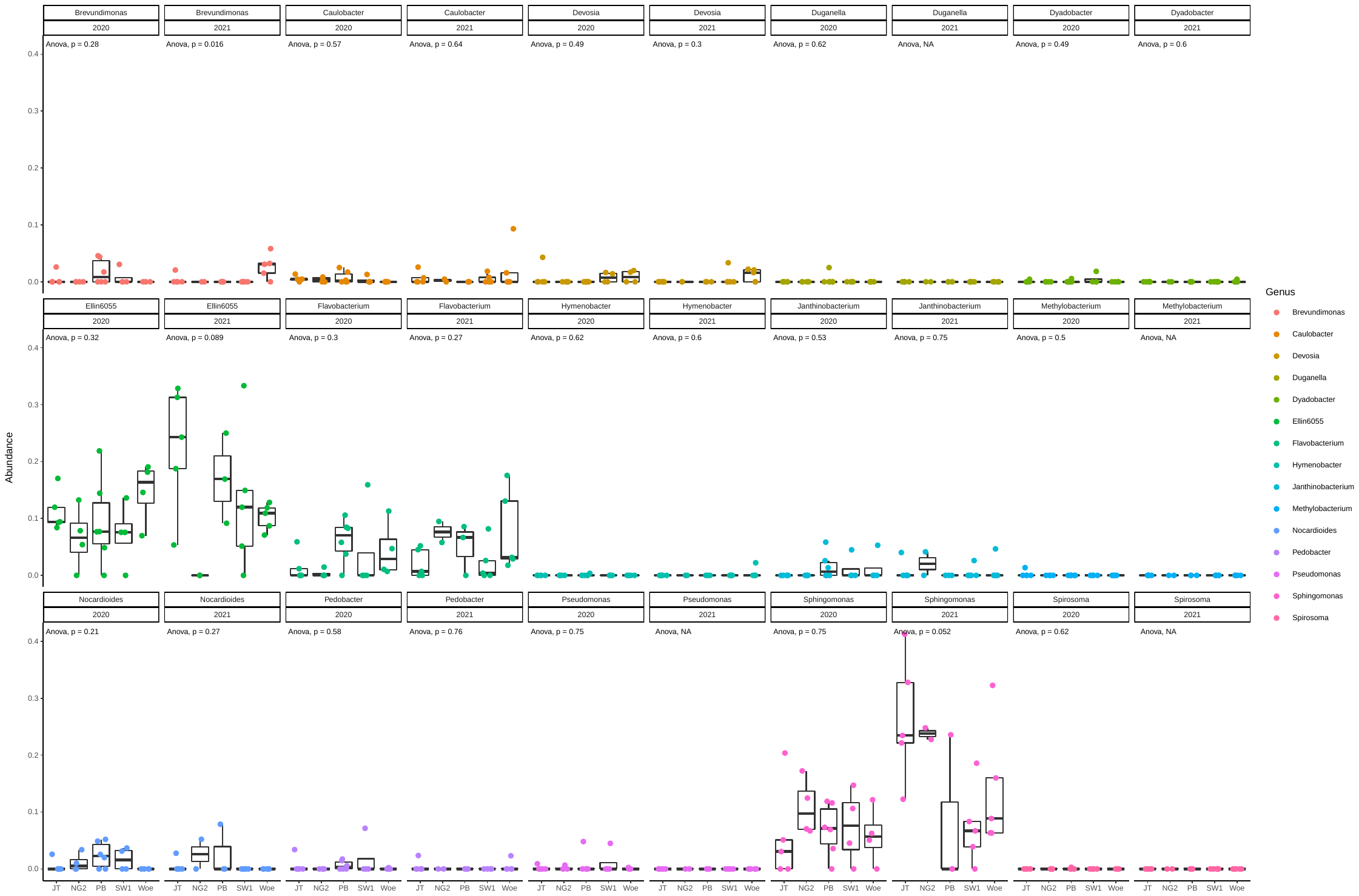

***Figure S8. Relative abundance of 15 of the 19 most prevalent core taxa that were observed at least once in soil samples, by location and year.***

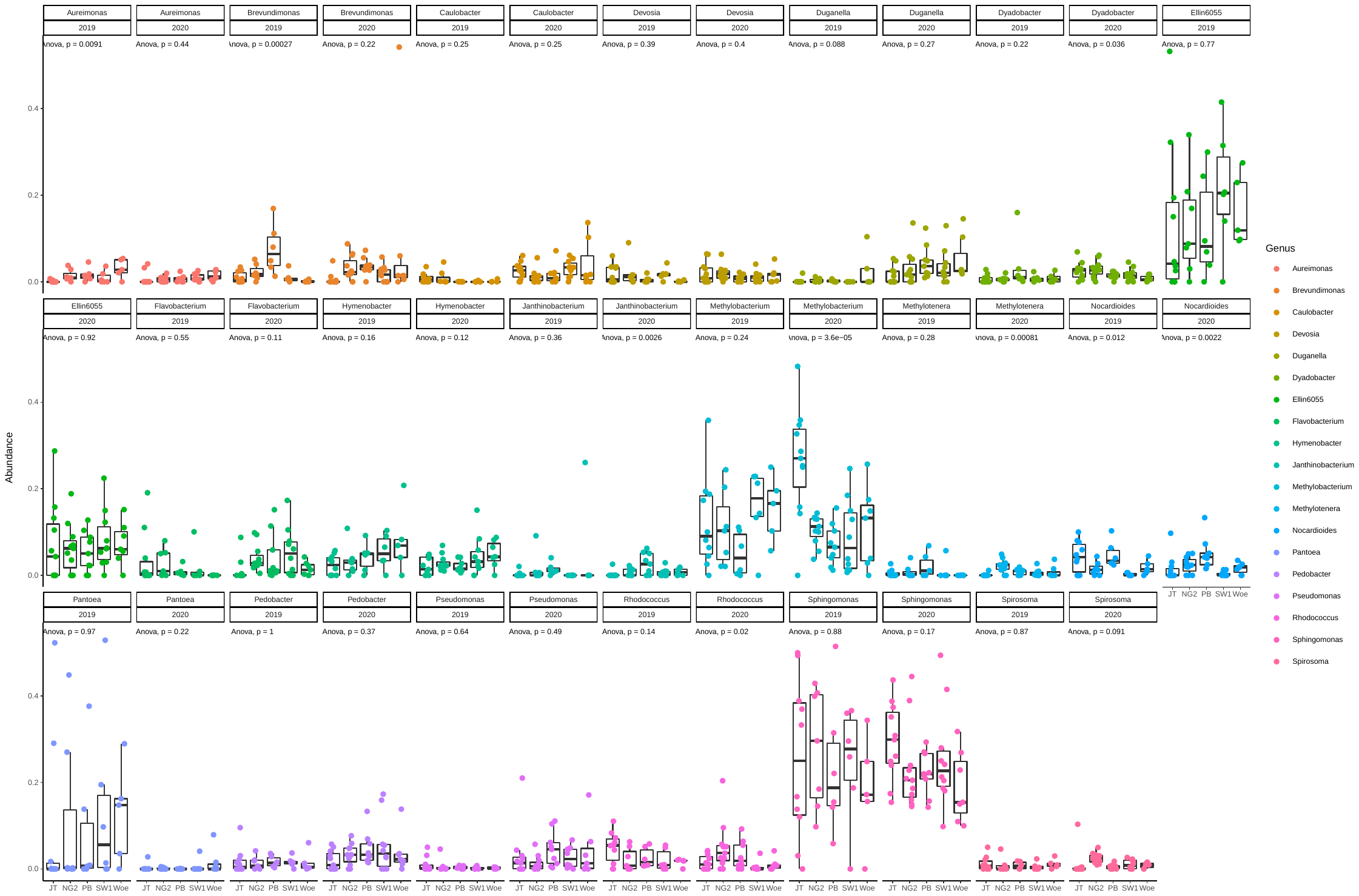

***Figure S9. Relative abundance the 19 most prevalent core taxa in A. thaliana, by year and location. P-values are ANOVA and show significant differences in abundance between locations.***

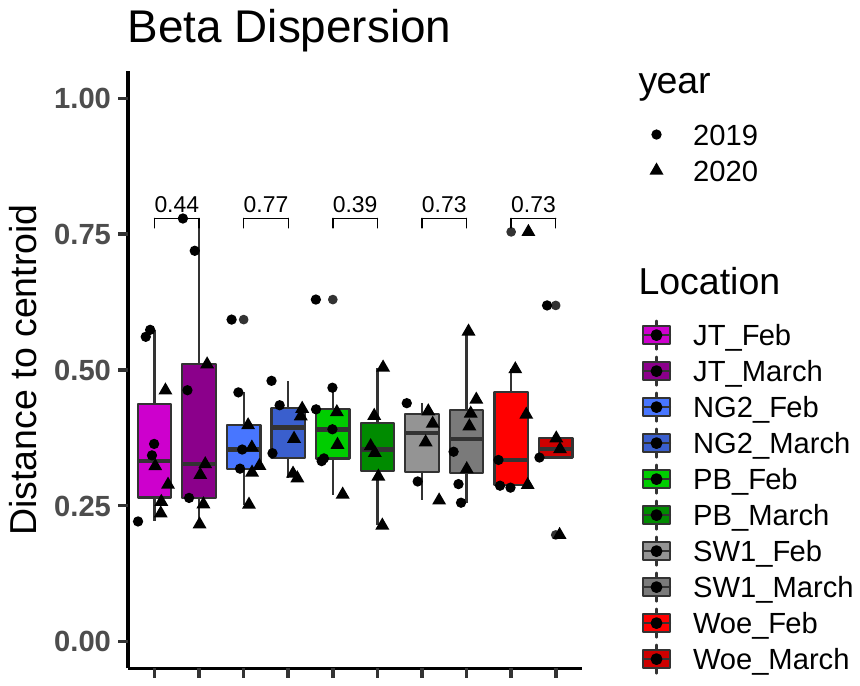

**Figure S10: Between-plant variance (betadispersion) calculated for each location individually looking at differences between the months.** P-values represent Wilcoxon rank-sum tests.

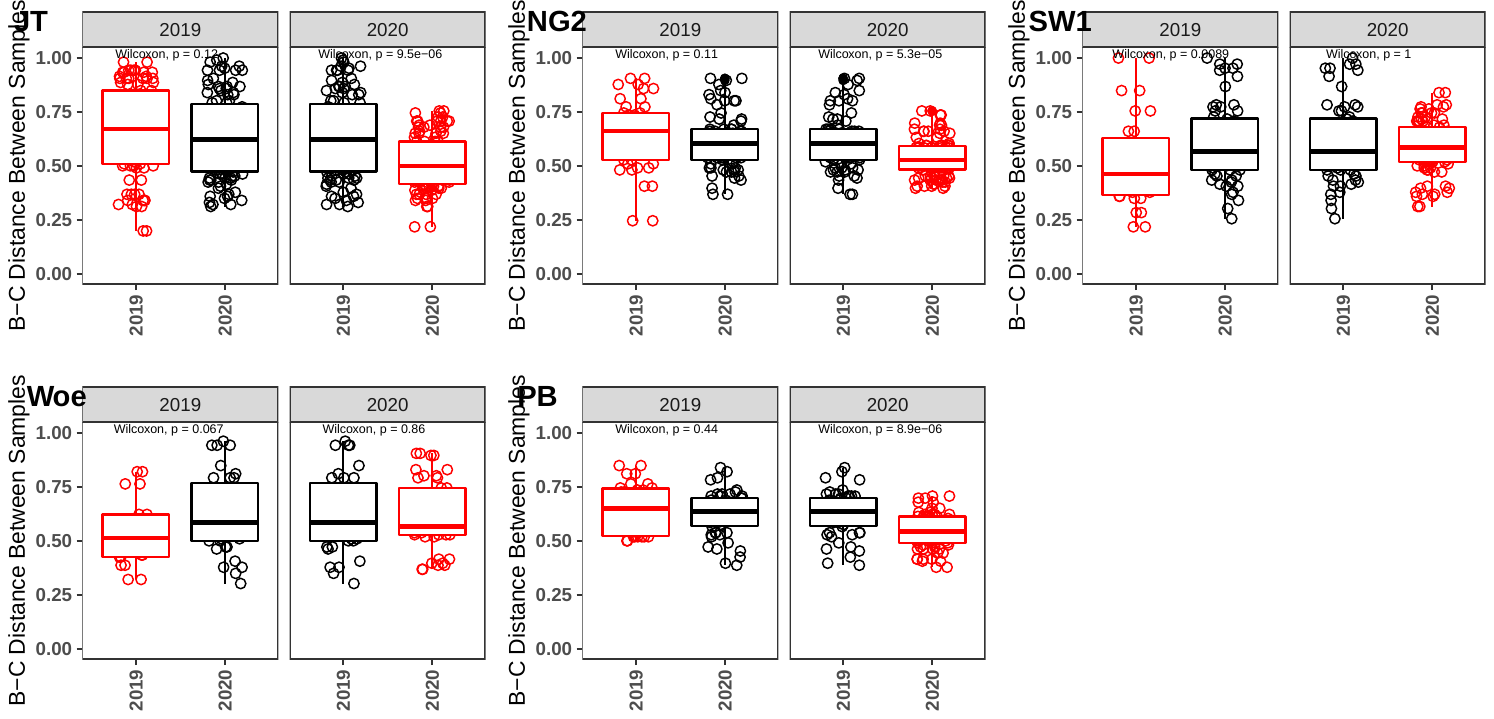

**Figure S11:** Bray-curtis distances between samples at each location comparing samples from the same year (red) vs samples from different years (black). Each plot compares samples of one year (plot header) to samples of the same or other year (x-axes). Samples were overall generally more similar within locations in 2020, except for a in increase at SW1.

**Figure S12 - Soil samples beta diversity**

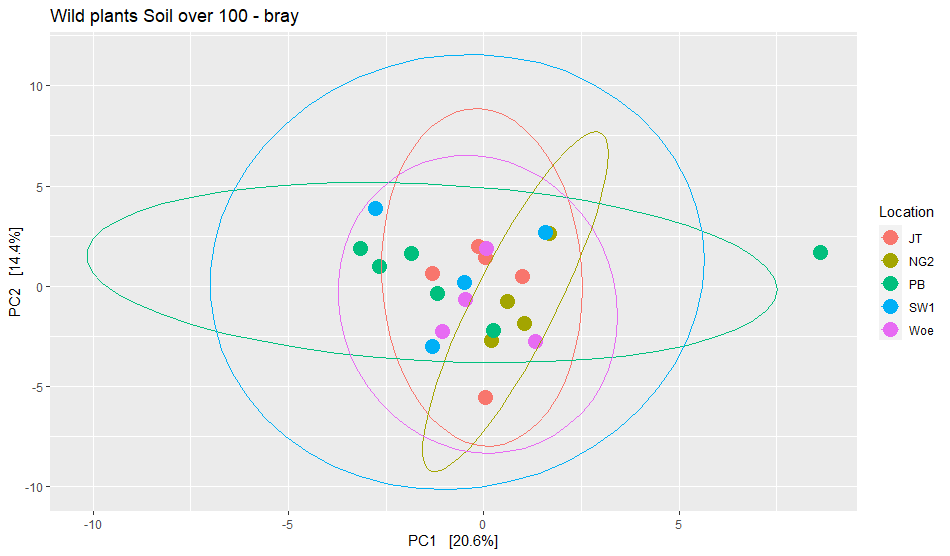

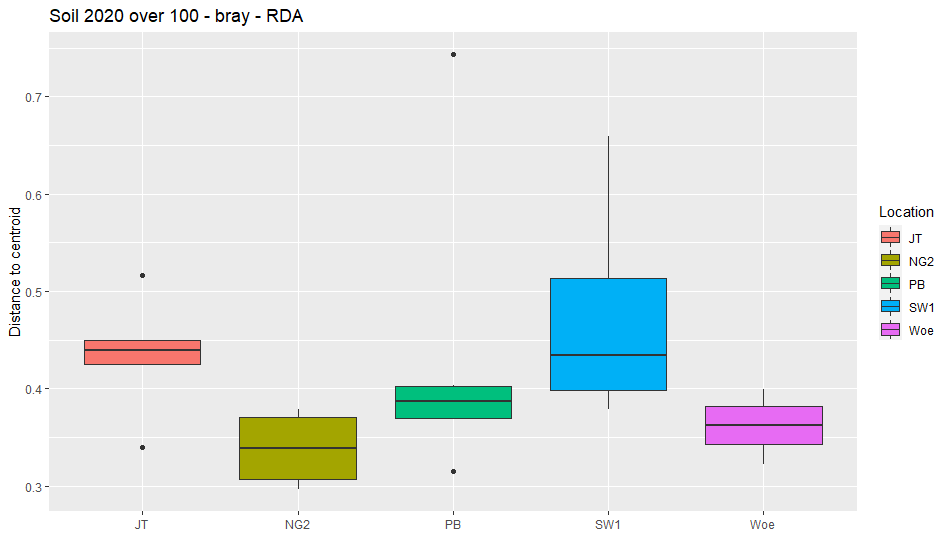

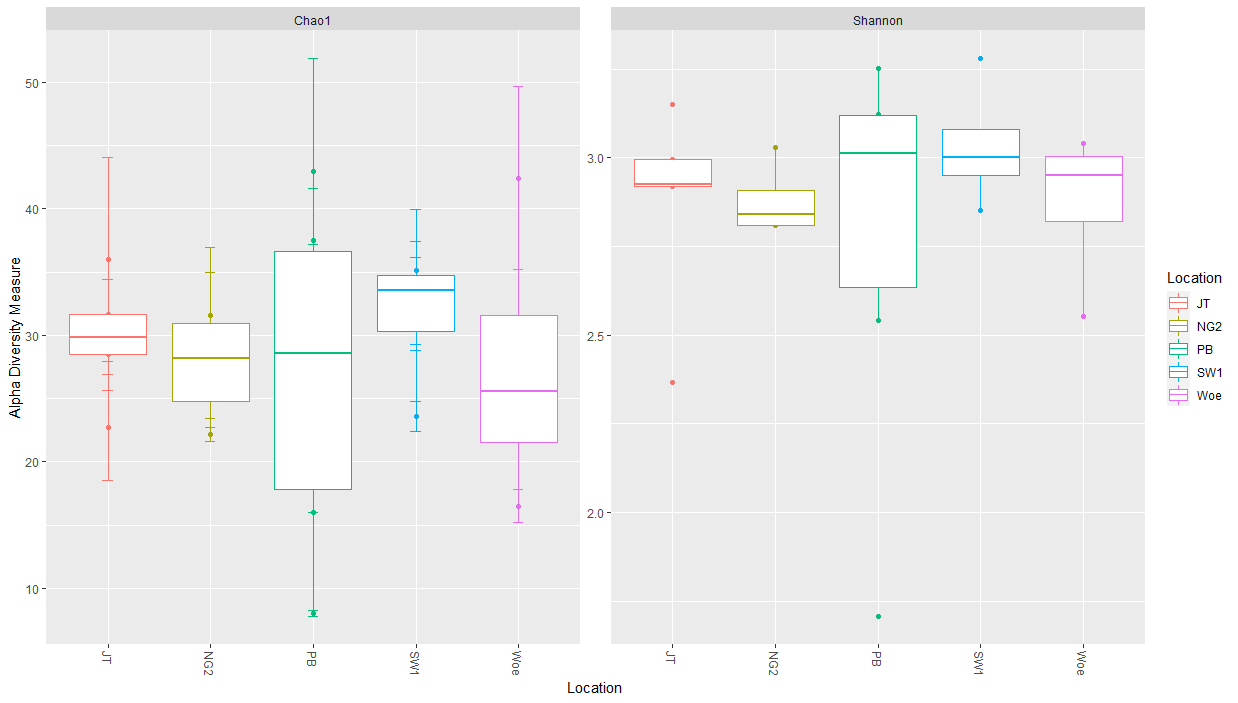

***Figure S12: The diversity of the bacterial communities of the soil samples was tested with different diversity measures (Statistics can be found in Supp Table 12) A****) Beta diversity differed significantly between the locations (R2=25.7%, p=0.003).* ***B****) Between-sample variation differs significantly between the locations NG2 and JT1.* ***C)*** *Alpha diversity is not significantly different between the locations.*

***A***

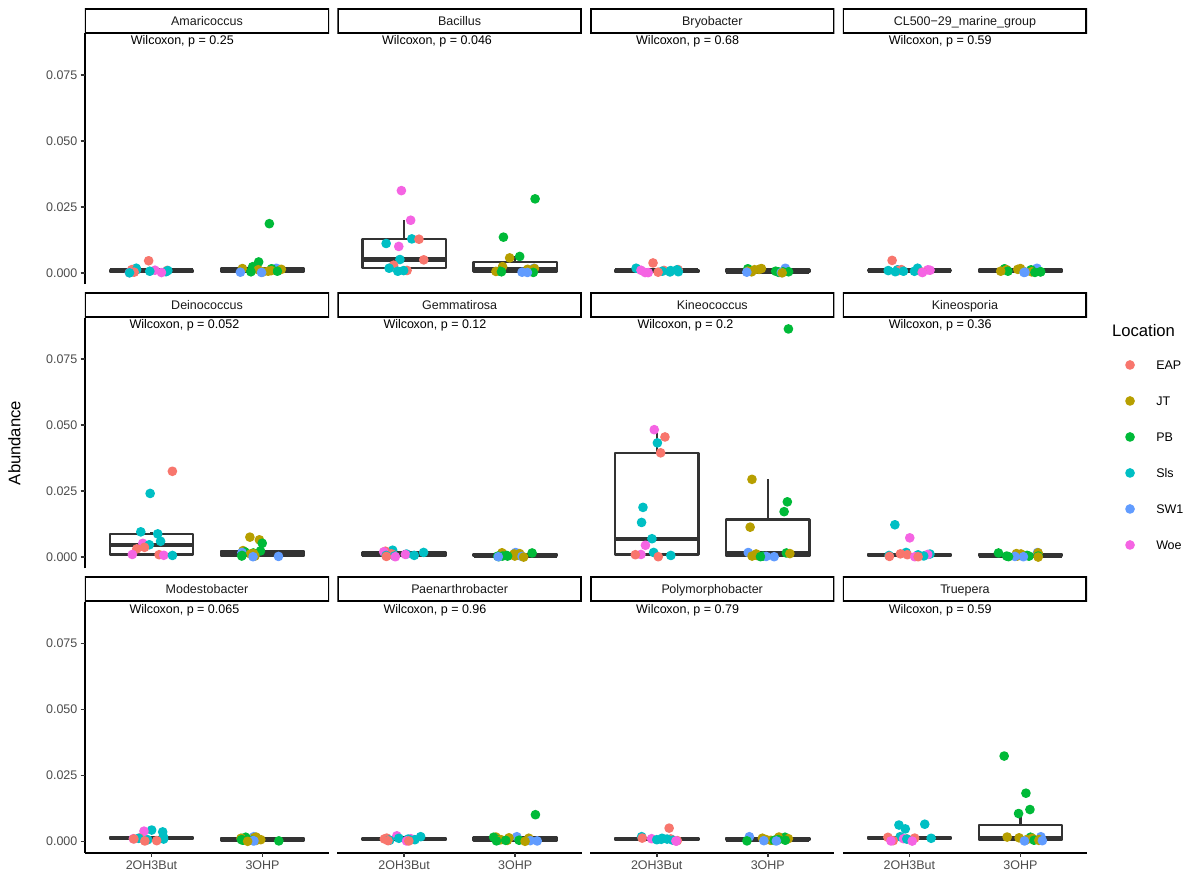

***B***

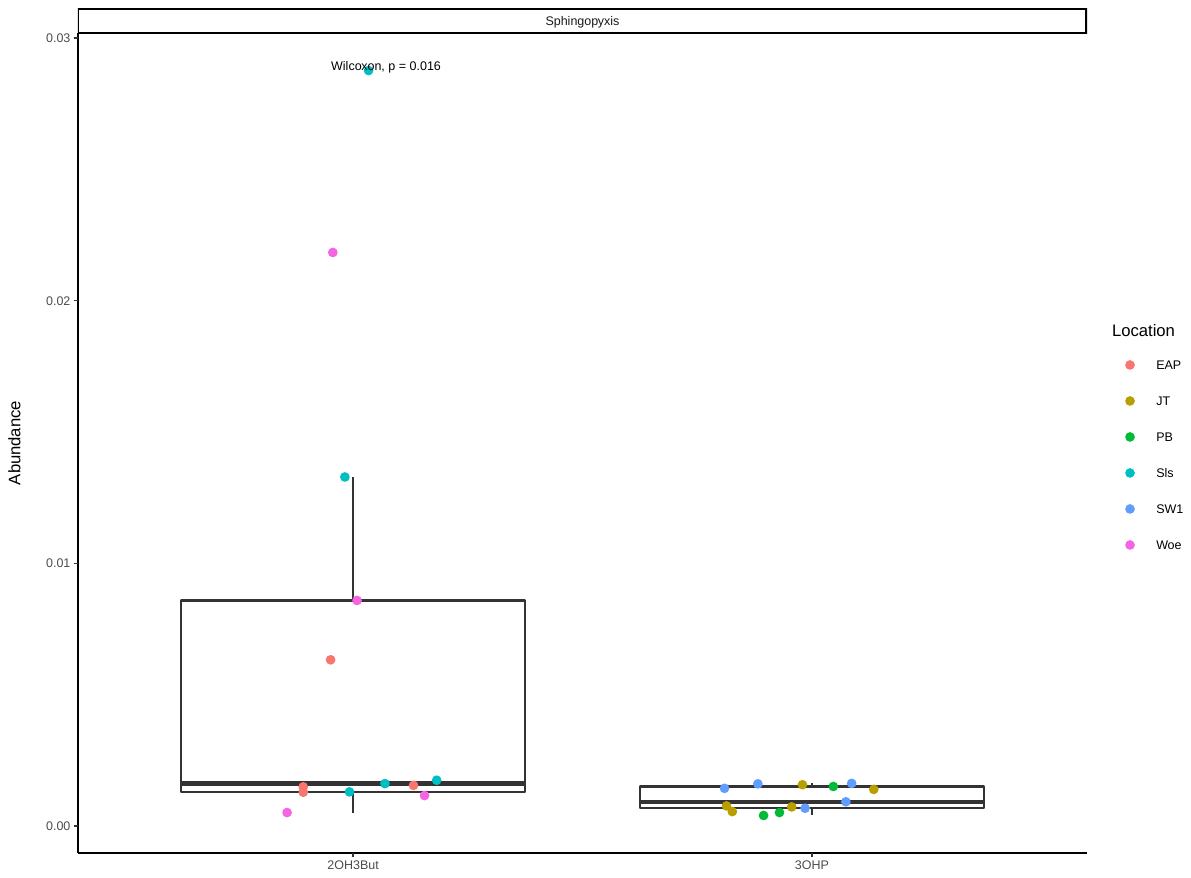

***Figure S13:* Taxa determined by a DESeq2 analysis to be enriched in one chemotype or the other (p<0.05). A) 2019 and B)2020.** Sample points are colored by location. Only *Bacillus* and *Sphingopyxis* were significantly enriched according to a Wilcoxon Rank-sum test (p<0.05).

**Supplementary Tables**

**Table S1: Primers used in this study**

| **Name** | **Primer sequence** | **primer length** | **Used for** | **References** |
| --- | --- | --- | --- | --- |
| nag59F | gcatctgtgttcactcgcc | 19 | Microsatellite diversity | Bell & Ecker 1994 |
| nag59R | ttaatacattagcccagacccg | 22 |  |  |
| nag111F | ctccagttggaagctaaaggg | 21 |  |  |
| nag111R | tgttttttaggacaaatggcg | 21 |  |  |
| nag158F | tcattttggccgacttagc | 19 |  |  |
| nag158R | acctgaaccatcctccgtc | 19 |  |  |
| 341-OH | TCCCTACACGACGCTCTTCCGATCTGACCTACGGGAGGCAGCAG | 44 | Amplicon Sequencing – Amplification Primer | Modified from Muyzer, Applied and Environomental Microbiology, 1999] / Caporaso et al., PNAS, 2010 |
| 799-OH | GGAGTTCAGACGTGTGCTCTTCCGATCTTGcmgggtatctaatcckgtt | 49 | Amplicon Sequencing – Amplification Primer | Modified from Chelius and Triplett, Microbial Ecology, 2001 / Bodenhausen et al., PlosONE, 2013 |
| Blc_16S-F5 | AACTTCTTTTCYHRGAGAAGAAR | 23 | Amplicon Sequencing – Blocking Oligo | Mayer et al. 2021 |
| Blc_16S-R1 | GCTTTCGCCRTTGGTGTTCYTTC | 23 | Amplicon Sequencing – Blocking Oligo | Mayer et al. 2021 |

**Table S2: Local *A. thaliana* plants sampled and used in this study.**

| **Population number** | **Name** | **Coordinates of Sampling Locations** | **Date found** | **Date collected** | **Micro-satellite Data** | **Flowering Time Data** | | **Gluco-sinolate Data** | **Analysis of bacterial community** | **Name in NASC database** |
| --- | --- | --- | --- | --- | --- | --- | --- | --- | --- | --- |
| 1a | NG1 | 50.9257433266571, 11.5828416046414 | Fall 2017 | 13.03.2018 | x | | x | x |  |  |
| 1b | NG2 |  | Fall 2017 | 09.05.2018 | x | | x | x | x | Je-1 |
| 2 | PB | 50.9255269205353, 11.585845678689 | Fall 2017 | 13.03.2018 | x | | x | x | x |  |
| 4 | Sls | 50.924174359466, 11.5725419221896 | 17.04.2018 | 17.04.2018 | x | | x | x | x |  |
| 5 | Woe | 50.904759003522, 11.5971417511712 | 05.04.2018 | 05.04.2018 | x | | x | x | x | Je-3 |
| 6a | SW1 | 50.919413031668, 11.5782925782252 | 12.04.2018 | 17.04.2018 | x | | x | x | x | Je-4 |
| 6b | SW2 | 50.919413031668, 11.5782925782252 | 12.04.2018 | 17.04.2018 | x | | x | x |  |  |
| 7a | JT1 | 50.930882676181, 11.58455821838 | 17.04.2018 | 17.04.2018 | x | |  | x | x | Je-2 |
| 7b | JT2 |  | 17.04.2018 | 17.04.2018 | x | |  | x |  |  |
| 9 | Loe | 50.945201672275, 11.6065010352618 | 20.04.2018 | 20.04.2018 | x | | x | x |  |  |
| 10 | Obi | 50.944425752363, 11.5992352752180 | 20.04.2018 | 20.04.2018 | x | | x |  | x |  |
| 11 | HB | 50.878036702216, 11.6178306950248 | 20.04.2018 | 20.04.2018 | x | |  |  | x |  |
| 12 | EAP | 50.9293679795480, 11.5826699432672 | 23.04.2018 | 23.04.2018 | x | | x | x | x |  |

**Table S3 – Correlation of beta diversity of *A. thaliana* and other plants to the factors plant species, location, and year.**

| **Arabidopsis/Other Plants \| 9 Locations** | | | | | | | | |
| --- | --- | --- | --- | --- | --- | --- | --- | --- |
| **Factor** | | | **Df** | **SumsOfSqs** | **MeanSqs** | **F.Model** | **R2** | **Pr(>F)** |
| **All Locations** |  |  |  |  |  |  |  |  |
| Year |  |  | 1 | 1.8978 | 1.8978 | 10.3495 | 0.0611 | 0.001 |
| Location |  |  | 8 | 3.1077 | 0.3885 | 2.1185 | 0.1001 | 0.001 |
| Plant |  |  | 1 | 0.5421 | 0.5421 | 2.9562 | 0.0175 | 0.004 |
| Year | Location |  | 8 | 2.6144 | 0.3268 | 1.7822 | 0.0842 | 0.001 |
| Year | Plant |  | 1 | 0.1806 | 0.1806 | 0.9850 | 0.0058 | 0.417 |
| Location | Plant |  | 8 | 1.7222 | 0.2153 | 1.1739 | 0.0555 | 0.136 |
| Year | Location | Plant | 8 | 1.9062 | 0.2383 | 1.2994 | 0.0614 | 0.037 |
| Residuals |  |  | 104 | 19.0708 | 0.1834 |  | 0.6144 |  |
| Total |  |  | 139 | 31.0419 |  |  | 1 |  |
| **NG2** | Year |  | 1 | 0.5888 | 0.5888 | 3.0295 | 0.1475 | 0.012 |
|  | Plant |  | 1 | 0.3209 | 0.3209 | 1.6509 | 0.0804 | 0.127 |
|  | Year | Plant | 1 | 0.1663 | 0.1663 | 0.8554 | 0.0417 | 0.523 |
|  | Residuals |  | 15 | 2.9155 | 0.1944 |  | 0.7304 |  |
|  | Total |  | 18 | 3.991 |  |  | 1 |  |
| **EAP** | Year |  | 1 | 0.45172788 | 0.45172788 | 2.3256305 | 0.16012954 | 0.026 |
|  | Plant |  | 1 | 0.32072321 | 0.32072321 | 1.65117921 | 0.1136907 | 0.123 |
|  | Year | Plant | 1 | 0.30041429 | 0.30041429 | 1.54662276 | 0.10649154 | 0.183 |
|  | Residuals |  | 9 | 1.74815 | 0.19423889 | NA | 0.61968822 | NA |
|  | Total |  | 12 | 2.82101538 | NA | NA | 1 | NA |
| **HKI** | Year |  | 1 | 0.58270667 | 0.58270667 | 3.77663822 | 0.18666684 | 0.018 |
|  | Plant |  | 1 | 0.54837413 | 0.54837413 | 3.55412219 | 0.1756686 | 0.008 |
|  | Year | Plant | 1 | 0.29334254 | 0.29334254 | 1.90121154 | 0.09397065 | 0.142 |
|  | Residuals |  | 11 | 1.69721667 | 0.15429242 | NA | 0.54369391 | NA |
|  | Total |  | 14 | 3.12164 | NA | NA | 1 | NA |
| **JT** | Year |  | 1 | 0.69392427 | 0.69392427 | 5.07100953 | 0.23327043 | 0.001 |
|  | Plant |  | 1 | 0.13367591 | 0.13367591 | 0.97686712 | 0.04493666 | 0.426 |
|  | Year | Plant | 1 | 0.09454131 | 0.09454131 | 0.69088217 | 0.03178112 | 0.724 |
|  | Residuals |  | 15 | 2.05262167 | 0.13684144 | NA | 0.6900118 | NA |
|  | Total |  | 18 | 2.97476316 | NA | NA | 1 | NA |
| **Obi** | Year |  | 1 | 0.37771519 | 0.37771519 | 2.40421283 | 0.16974379 | 0.014 |
|  | Plant |  | 1 | 0.16469435 | 0.16469435 | 1.04830381 | 0.07401302 | 0.407 |
|  | Year | Plant | 1 | 0.26884815 | 0.26884815 | 1.71125806 | 0.12081935 | 0.085 |
|  | Residuals |  | 9 | 1.41395 | 0.15710556 | NA | 0.63542383 | NA |
|  | Total |  | 12 | 2.22520769 | NA | NA | 1 | NA |
| **PB** | Year |  | 1 | 0.32177078 | 0.32177078 | 1.68829014 | 0.10411629 | 0.075 |
|  | Plant |  | 1 | 0.10863524 | 0.10863524 | 0.5699952 | 0.03515141 | 0.865 |
|  | Year | Plant | 1 | 0.18242143 | 0.18242143 | 0.9571419 | 0.05902662 | 0.454 |
|  | Residuals |  | 13 | 2.47766667 | 0.19058974 | NA | 0.80170567 | NA |
|  | Total |  | 16 | 3.09049412 | NA | NA | 1 | NA |
| **Sls** | Year |  | 1 | 0.50220923 | 0.50220923 | 2.06957668 | 0.1304806 | 0.036 |
|  | Plant |  | 1 | 0.20172063 | 0.20172063 | 0.83127967 | 0.05240969 | 0.607 |
|  | Year | Plant | 1 | 0.23303556 | 0.23303556 | 0.96032675 | 0.06054572 | 0.444 |
|  | Residuals |  | 12 | 2.91195333 | 0.24266278 | NA | 0.75656399 | NA |
|  | Total |  | 15 | 3.84891875 | NA | NA | 1 | NA |
| **SW1** | Year |  | 1 | 0.47736417 | 0.47736417 | 2.71380407 | 0.15388978 | 0.008 |
|  | Plant |  | 1 | 0.3129 | 0.3129 | 1.77882915 | 0.10087081 | 0.079 |
|  | Year | Plant | 1 | 0.20089667 | 0.20089667 | 1.14209283 | 0.06476385 | 0.318 |
|  | Residuals |  | 12 | 2.11082667 | 0.17590222 | NA | 0.68047556 | NA |
|  | Total |  | 15 | 3.1019875 | NA | NA | 1 | NA |
| **Woe** | Year |  | 1 | 0.34873429 | 0.34873429 | 1.60067757 | 0.1335635 | 0.132 |
|  | Plant |  | 1 | 0.17285689 | 0.17285689 | 0.79340678 | 0.06620333 | 0.577 |
|  | Year | Plant | 1 | 0.34647549 | 0.34647549 | 1.59030978 | 0.13269839 | 0.126 |
|  | Residuals |  | 8 | 1.74293333 | 0.21786667 | NA | 0.66753479 | NA |
|  | Total |  | 11 | 2.611 | NA | NA | 1 | NA |

**Table S4: Abundance of leaf bacteria at the phyla level in absolute and relative amount, separated by populations**

|  | **Arabidopsis plants** | | | | | | | | | |
| --- | --- | --- | --- | --- | --- | --- | --- | --- | --- | --- |
|  | **Absolute read counts** | | | | | **Relative abundance** | | | | |
| **Phylum** | **JT1** | **NG2** | **PB** | **SW1** | **Woe** | **JT** | **NG2** | **PB** | **SW1** | **Woe** |
| Proteobacteria | 50554 | 31662 | 28169 | 39416 | 33294 | 0.653 | 0.721 | 0.704 | 0.631 | 0.775 |
| Bacteroidetes | 14609 | 5663 | 6048 | 21081 | 5252 | 0.189 | 0.129 | 0.151 | 0.337 | 0.122 |
| Actinobacteria | 11864 | 6247 | 5115 | 1926 | 3928 | 0.153 | 0.142 | 0.128 | 0.031 | 0.091 |
| Deinococcus-Thermus | 258 | 199 | 612 | 9 | 87 | 0.003 | 0.005 | 0.015 | 0.000 | 0.002 |
| Firmicutes | 92 | 125 | 78 | 31 | 407 | 0.001 | 0.003 | 0.002 | 0.000 | 0.009 |
| Gemmatimonadetes | 36 | 43 | 6 | 23 | 4 | 0.001 | 0.004 | 0.001 | 0.001 | 0.000 |

**Table S5: Core taxa determined independently for each location or each year.** The genus names in bold are shared across locations and years. Genus names in bold are included in the 19 taxa shown in Figure 3 (was among the top 10 most prevalent at least at one location)

|  |  | **Location** | | | | |
| --- | --- | --- | --- | --- | --- | --- |
| **2019** | **2020** | **JT** | **NG2** | **PB** | **SW1** | **Woe** |
| **Sphingomonas** | **Sphingomonas** | **Sphingomonas** | **Sphingomonas** | **Sphingomonas** | **Sphingomonas** | **Sphingomonas** |
| **Ellin6055** | **Ellin6055** | **Ellin6055** | **Ellin6055** | **Ellin6055** | **Ellin6055** | **Ellin6055** |
| **Methylobacterium** | **Methylobacterium** | **Methylobacterium** | **Methylobacterium** | **Methylobacterium** | **Methylobacterium** | **Methylobacterium** |
| Pantoea |  |  |  |  |  | Pantoea |
| **Rhodococcus** | **Rhodococcus** | **Rhodococcus** | **Rhodococcus** | **Rhodococcus** | **Rhodococcus** | **Rhodococcus** |
| Brevundimonas | Brevundimonas |  | Brevundimonas | Brevundimonas | Brevundimonas | Brevundimonas |
| Aureimonas | Aureimonas |  | Aureimonas | Aureimonas | Aureimonas | Aureimonas |
| **Caulobacter** | **Caulobacter** | **Caulobacter** | **Caulobacter** | **Caulobacter** | **Caulobacter** | **Caulobacter** |
| **Pedobacter** | **Pedobacter** | **Pedobacter** | **Pedobacter** | **Pedobacter** | **Pedobacter** | **Pedobacter** |
| Flavobacterium | Flavobacterium |  | Flavobacterium | Flavobacterium | Flavobacterium | Flavobacterium |
|  |  |  |  | Kineococcus |  | Kineococcus |
| **Devosia** | **Devosia** | **Devosia** | **Devosia** | **Devosia** | **Devosia** | **Devosia** |
| **Pseudomonas** | **Pseudomonas** | **Pseudomonas** | **Pseudomonas** | **Pseudomonas** | **Pseudomonas** | **Pseudomonas** |
|  | Janthinobacterium |  | Janthinobacterium | Janthinobacterium |  | Janthinobacterium |
| **Hymenobacter** | **Hymenobacter** | **Hymenobacter** | **Hymenobacter** | **Hymenobacter** | **Hymenobacter** | **Hymenobacter** |
| Nocardioides | Nocardioides | Nocardioides | Nocardioides | Nocardioides |  | Nocardioides |
|  | Duganella |  | Duganella | Duganella | Duganella | Duganella |
| Variovorax | Variovorax | Variovorax | Variovorax |  | Variovorax | Variovorax |
| Bacillus |  |  |  |  |  |  |
|  | Ralstonia | Ralstonia |  | Ralstonia | Ralstonia |  |
|  |  |  |  | Rathayibacter |  | Rathayibacter |
|  | Nakamurella |  | Nakamurella | Nakamurella |  | Nakamurella |
|  | Mycobacterium |  | Mycobacterium |  | Mycobacterium | Mycobacterium |
|  |  |  |  |  |  | Actinoplanes |
|  |  |  |  | Truepera |  |  |
| **Dyadobacter** | **Dyadobacter** | **Dyadobacter** | **Dyadobacter** | **Dyadobacter** | **Dyadobacter** | **Dyadobacter** |
| Methylotenera | Methylotenera |  | Methylotenera | Methylotenera | Methylotenera | Methylotenera |
|  |  |  | Bosea |  |  |  |
|  | Sediminibacterium |  |  | Craurococcus |  |  |
|  |  |  |  |  | Rhizobacter |  |
|  |  |  |  |  | Sediminibacterium | Aeromicrobium |
|  |  |  |  |  |  | Deinococcus |
| Massilia | Massilia |  | Massilia | Massilia | Massilia | Massilia |
| Patulibacter |  |  |  |  |  | Patulibacter |
| **Spirosoma** | **Spirosoma** | **Spirosoma** | **Spirosoma** | **Spirosoma** | **Spirosoma** | **Spirosoma** |
| Luteimonas | Luteimonas |  | Luteimonas |  | Luteimonas |  |
|  | Bdellovibrio |  |  |  | Bdellovibrio | Bdellovibrio |
|  |  |  |  |  |  | Roseomonas |
|  |  |  |  | Rubellimicrobium |  | Rubellimicrobium |
|  |  |  |  | Amaricoccus |  | Actinomycetospora |
|  |  |  |  | Ilumatobacter |  |  |
|  |  |  |  | Aquamicrobium |  |  |
|  |  |  | Hansschlegelia |  |  |  |
|  |  |  | Acidovorax |  |  |  |
|  |  |  | Fibrella |  |  |  |
|  |  |  | Fibrisoma |  | Fibrisoma |  |
|  |  |  | Rhodopseudomonas |  |  |  |
|  |  |  | Ferruginibacter |  |  |  |
|  |  |  | Pseudorhodobacter |  |  |  |

**Table S6: Abundances of the different phyla in the soil samples in absolute and relative amount, separated by populations**

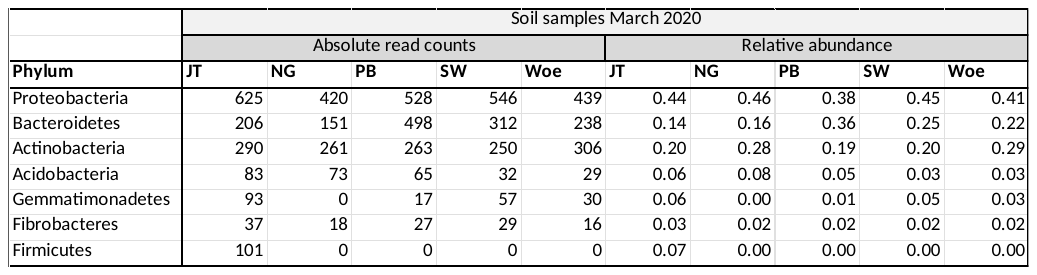

**Table S7 - Adonis values calculated for the beta diversity comparisons of the *Arabidopsis* plants for the three variables location, month and year.**

| **Arabidopsis** | | | | | | | | |
| --- | --- | --- | --- | --- | --- | --- | --- | --- |
| **Factor** | | | **Df** | **SumsOfSqs** | **MeanSqs** | **F.Model** | **R2** | **Pr(>F)** |
| Location |  |  | 4 | 1.5176 | 0.3794 | 2.4055 | 0.09316 | 0.001 |
| Month |  |  | 1 | 0.3419 | 0.34189 | 2.1677 | 0.02099 | 0.019 |
| Year |  |  | 1 | 0.9347 | 0.93468 | 5.9261 | 0.05738 | 0.001 |
| Location | Month |  | 4 | 1.0582 | 0.26454 | 1.6772 | 0.06496 | 0.007 |
| Location | Year |  | 4 | 0.8128 | 0.2032 | 1.2883 | 0.0499 | 0.113 |
| Month | Year |  | 1 | 0.1697 | 0.16974 | 1.0762 | 0.01042 | 0.345 |
| Location | Month | Year | 3 | 1.5184 | 0.50612 | 3.2089 | 0.09321 | 0.001 |
| Residuals |  |  | 63 | 9.9366 | 0.15772 |  | 0.60999 |  |
| Total |  |  | 81 | 16.2898 |  |  | 1 |  |

**Table S8 – Adonis values calculated for the beta diversity comparisons of the *Arabidopsis* plants of the individual locations focusing on year and month**

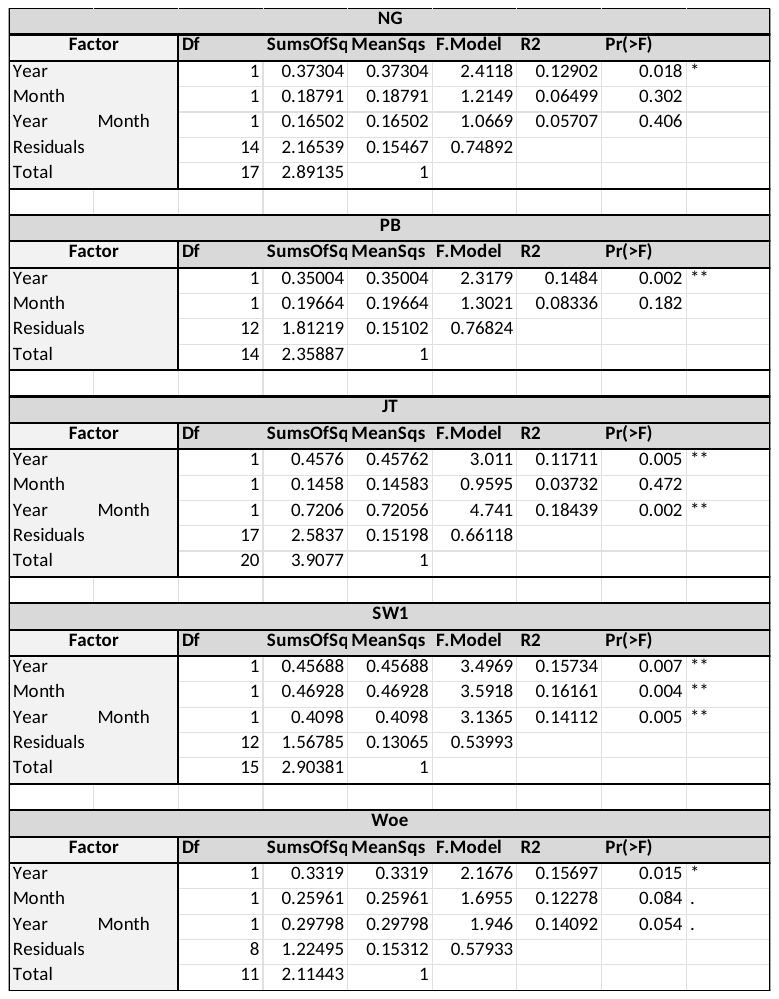

**Table S9 – DeSeq2 on individual locations to identify additional taxa (apart form the core taxa) that were significantly different between the locations**

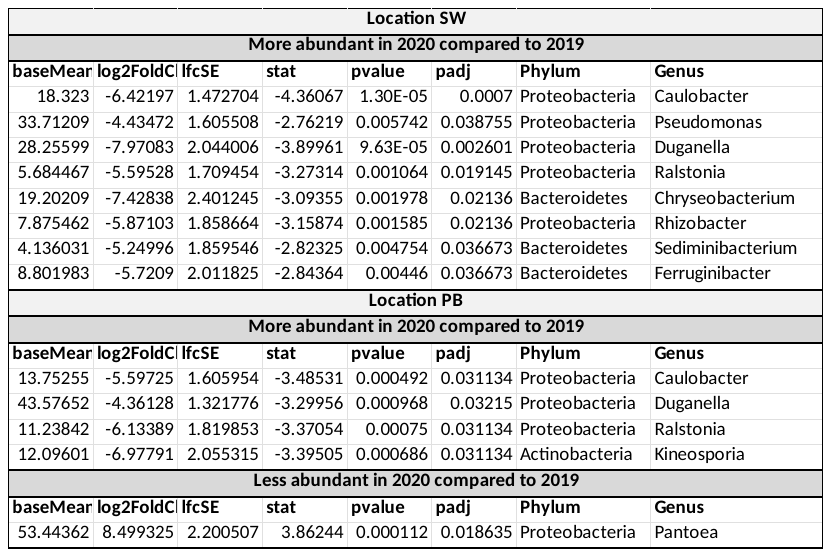

**Table S10 – Adonis (permanova) values calculated for the beta diversity comparisons of the *Arabidopsis* plants focusing on location and month.**

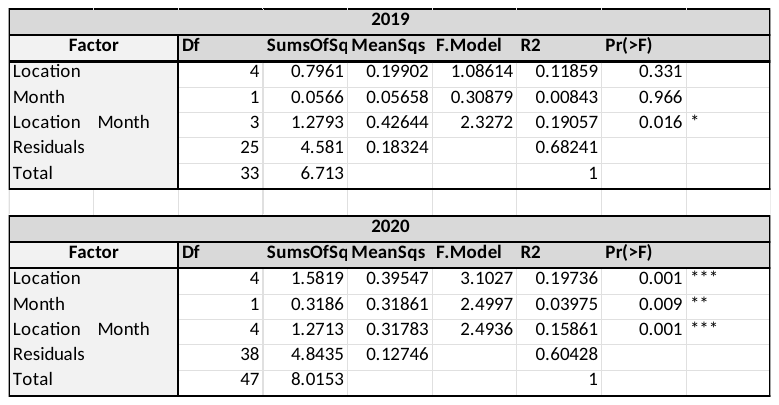

**Table S11 –**

**Anova values calculated for the alpha diversity of the *Arabidopsis* plants of the individual locations focusing on the differences between the years**

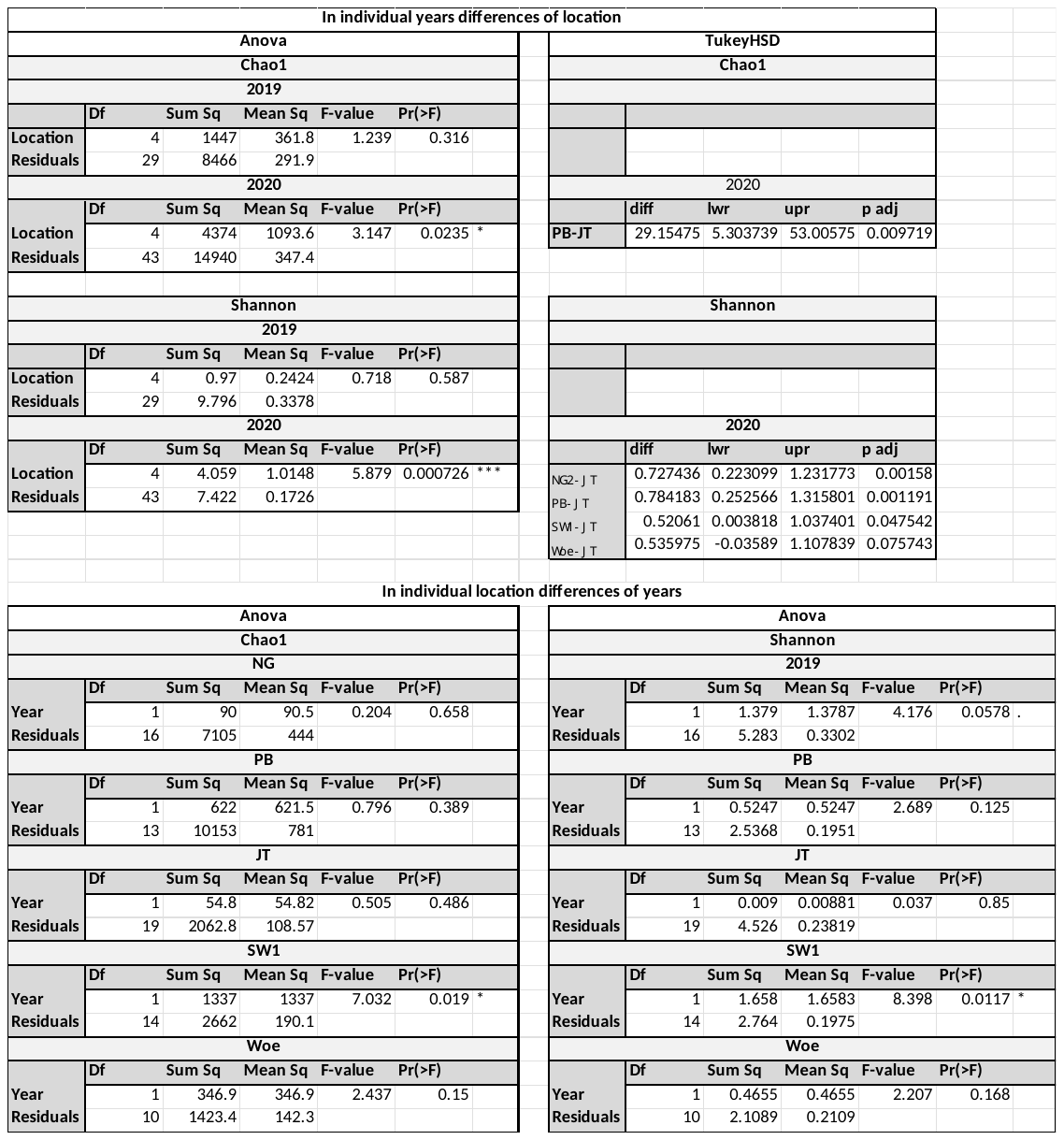

**Table S12 - DeSeq2 on individual years to identify additional taxa (apart form the core taxa) that were significantly different between the locations**

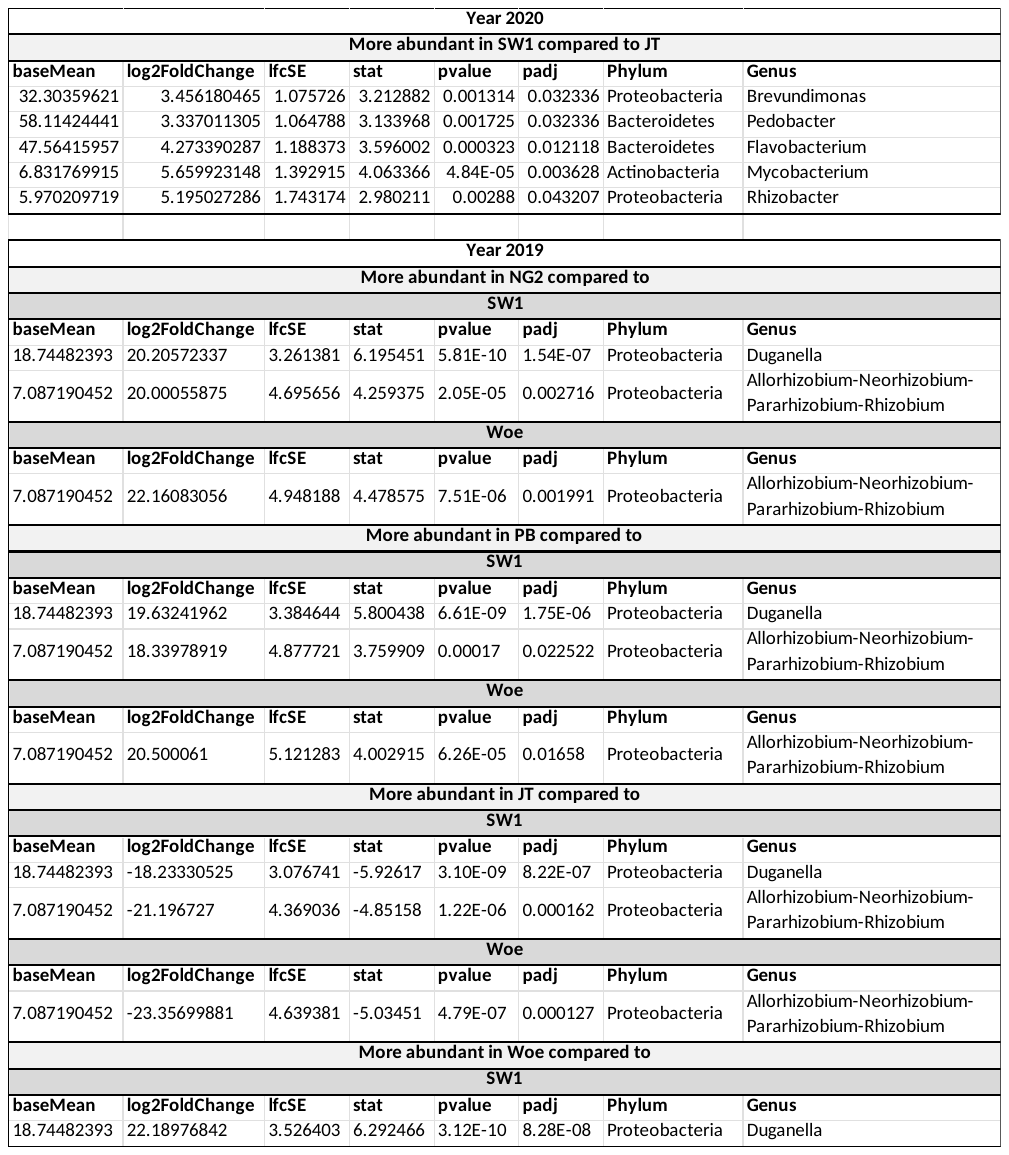

**Table S13: Overview of diversity stats of soil samples**

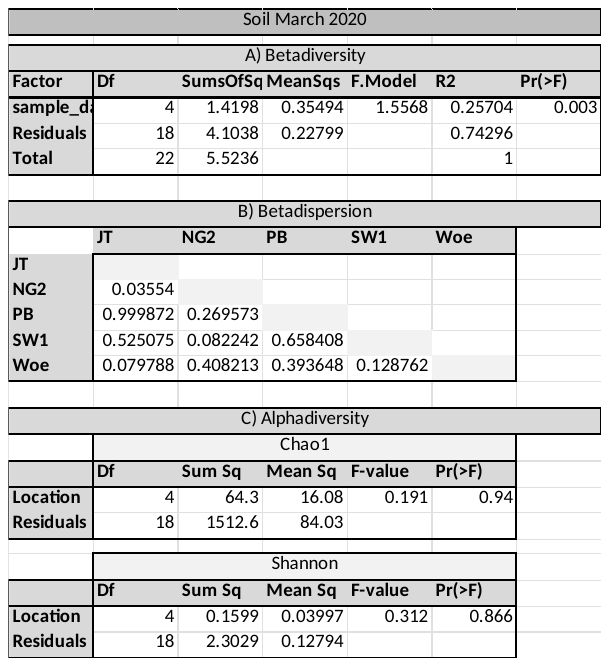
